# Genomic and phenotypic heterogeneity of clinical isolates of the human pathogens *Aspergillus fumigatus, Aspergillus lentulus* and *Aspergillus fumigatiaffinis*

**DOI:** 10.1101/2020.02.28.970384

**Authors:** Renato Augusto Corrêa dos Santos, Jacob L. Steenwyk, Olga Rivero-Menendez, Matthew E. Mead, Lilian Pereira Silva, Rafael Wesley Bastos, Ana Alastruey-Izquierdo, Gustavo Henrique Goldman, Antonis Rokas

## Abstract

Fungal pathogens are a global threat to human health. For example, fungi from the genus *Aspergillus* cause a spectrum of diseases collectively known as aspergillosis. Most of the >200,000 life-threatening aspergillosis infections per year worldwide are caused by *Aspergillus fumigatus*. Recently, molecular typing techniques have revealed that aspergillosis can also be caused by organisms that are phenotypically similar to *A. fumigatus* but genetically distinct, such as *Aspergillus lentulus* and *Aspergillus fumigatiaffinis*. Importantly, some of these so-called cryptic species exhibit different virulence and drug susceptibility profiles than *A. fumigatus*, however, our understanding of their biology and pathogenic potential has been stymied by the lack of genome sequences and phenotypic profiling. To fill this gap, we phenotypically characterized the virulence and drug susceptibility of 15 clinical strains of *A. fumigatus, A. lentulus*, and *A. fumigatiaffinis* from Spain and sequenced their genomes. We found heterogeneity in virulence and drug susceptibility across species and strains. Genes known to influence drug susceptibility (*cyp51A* and *fks1*) vary in paralog number and sequence among these species and strains and correlate with differences in drug susceptibility. Similarly, genes known to be important for virulence in *A. fumigatus* showed variability in number of paralogs across strains and across species. Characterization of the genomic similarities and differences of clinical strains of *A. lentulus, A. fumigatiaffinis, and A. fumigatus* that vary in disease-relevant traits will advance our understanding of the variance in pathogenicity between *Aspergillus* species and strains that are collectively responsible for the vast majority of aspergillosis infections in humans.

## INTRODUCTION

Aspergillosis is a major health problem, with rapidly evolving epidemiology and new groups of at-risk patients (Patterson et al., 2016). Aspergillosis infections are usually caused by inhalation of airborne asexual spores (conidia) of *Aspergillus fumigatus* and a few other *Aspergillus* species (Rokas et al., 2020). Aspergillosis covers a spectrum of diseases (Latgé and Chamilos, 2020). For example, non-invasive diseases caused by *Aspergillus*, such as aspergilloma, are currently classified as chronic pulmonary aspergillosis and are commonly associated to pulmonary tuberculosis (Denning et al., 2016). In atopic patients, the most severe form of aspergillosis is allergic bronchopulmonary aspergillosis (ABPA), which develops following sensibilization to *A. fumigatus* allergens in atopic patients with cystic fibrosis or individuals with genetic predisposition to ABPA (Agarwal et al., 2013). However, the most common invasive type of infection is invasive pulmonary aspergillosis (IPA), whose risk is significantly increased in immunocompromised individuals, in patients with acute leukaemia and recipients of hematopoietic stem cells transplantation, or in solid-organ transplant recipients (Brown et al., 2012). Importantly, IPA has recently been described in new groups of traditionally low-risk patients, such as patients in intensive care units recovering from bacterial sepsis (Latgé and Chamilos, 2020).

Although *A. fumigatus* is the major etiologic agent of aspergillosis, a few other *Aspergillus* species, such as *Aspergillus flavus, Aspergillus terreus, Aspergillus niger*, and *Aspergillus nidulans*, can also cause infections (Zakaria et al., 2020). While most of these pathogens can be phenotypically easily distinguished, infections can also be caused by *Aspergillus* species that are morphologically very similar to *A. fumigatus* (Rokas et al., 2020). These close pathogenic relatives of *A. fumigatus* are considered sibling species or cryptic species because they are undistinguishable from each other and from *A. fumigatus* by classical identification methods (Alastruey-Izquierdo et al., 2014); these species vary mostly in their colony growth, robustness of the production of conidia, conidial surface markings, presence and absence of septation in phialides, and maximum growth temperatures (Taylor et al., 2000; Balajee et al., 2005; Katz et al., 2005). As a result of their near identical morphological characteristics, most of these cryptic species have only recently been described. For example, *Aspergillus lentulus* was first described in 2005 in a case of human aspergillosis (Balajee et al., 2005). Similarly, *A. fumigatiaffinis*, another pathogenic species that is closely related to *A. fumigatus*, was first described in 2005 (Hong et al., 2005). Even though cryptic species were only discovered relatively recently, understanding their genetic and phenotypic similarities and differences from the major pathogen *A. fumigatus* is important for two reasons. First, their prevalence in the clinic has been estimated to be between 11 and 19% (Balajee et al., 2009; Alastruey-Izquierdo et al., 2014; Negri et al., 2014). Second, several of these species, including *A. lentulus* and *A. fumigatiaffinis*, have been shown to differ in their drug susceptibility to amphotericin B and azoles compared to *A. fumigatus* (Alastruey-Izquierdo et al., 2014).

An emerging realization in the study of fungal pathogens is the presence of phenotypic heterogeneity among strains of the same species, especially with respect to virulence and antifungal drug susceptibility (Keller, 2017). For example, a recent study reported strong correlation between *in vitro* hypoxic growth phenotypes and virulence among *A. fumigatus* strains (Kowalski et al., 2016). Similarly, *A. fumigatus* strains have previously been show to exhibit great quantitative and qualitative heterogeneity in light response (Fuller et al., 2016); in this case, heterogeneity in light response was not associated with heterogeneity in virulence. Finally, Ries *et al*. found a high heterogeneity among *A. fumigatus* strains with regard to nitrogen acquisition and metabolism during infection and correlation between nitrogen catabolite repression-related protease secretion and virulence (Ries et al., 2019). These studies highlight the biological and clinical relevance of understanding strain heterogeneity in *Aspergillus* pathogens. However, comparisons of strain heterogeneity in virulence and drug resistance profiles among clinical strains in *A. fumigatus* and closely related cryptic species, such as *A. lentulus* and *A. fumigatiaffinis*, are lacking.

To address this gap in the field, we phenotypically characterized and sequenced the genomes of 15 clinical strains of *A. fumigatus, A. lentulus*, and *A. fumigatiaffinis* from Spain. We found strain heterogeneity in both virulence and drug susceptibility profiles within each species as well as differences between the three species. We found that genes known to influence drug susceptibility, such as *cyp51A*, exhibit variation in their numbers of paralogs and sequence among these species and strains. Similarly, we found variability in the number of paralogs within and between species in many genes known to be important for virulence in *A. fumigatus*. Characterization of the genomic similarities and differences of clinical strains of *A. lentulus, A. fumigatiaffinis, and A. fumigatus* that vary in disease-relevant traits will advance our understanding of the variation in pathogenicity between *Aspergillus* species and strains that are collectively responsible for the vast majority of aspergillosis infections in humans.

## MATERIALS AND METHODS

### Strains and species identification

To understand the degree of genomic heterogeneity among strains, we sequenced six clinical strains of *A. fumigatus*, five of *A. lentulus* and four of *A. fumigatiaffinis* available in the Mycology Reference Laboratory of the National Center for Microbiology (CNM) in Instituto de Salud Carlos III in Spain (Supplementary Table 1). For initial species identification, we sequenced the Internal Transcribed Spacer region (ITS) and beta-tubulin (*benA*) gene amplicons (primer pairs in Supplementary Table 2). We downloaded reference sequences for the type strains of *A. fumigatiaffinis* IBT12703 and *A. lentulus* IFM54703, and of *Aspergillus clavatus* NRRL1 (section *Clavati*), which we used as the outgroup. We aligned DNA sequences with MAFFT v.7.397 (Katoh and Standley, 2013), followed by model selection and phylogenetic inference in IQ-TREE v.1.6.7 (Nguyen et al., 2015).

### Characterization of virulence and antifungal susceptibility profiles

To understand the pathogenic potential of the 15 clinical strains, we carried out virulence assays using the moth *Galleria mellonella* model of fungal disease (Fuchs et al., 2010). Briefly, we obtained moth larvae by breeding adult moths that were kept for 24 hr prior to infection under starvation, in the dark, and at a temperature of 37 °C. We selected only larvae that were in the sixth and final stage of larval development. We harvested fresh asexual spores (conidia) from each strain from yeast extract-agar-glucose (YAG) plates in PBS solution and filtered through a Miracloth (Calbiochem). For each strain, we counted the spores using a hemocytometer and created a 2 × 10^8^ conidia/ml stock suspension. We determined the viability of the administered inoculum by plating a serial dilution of the conidia on YAG medium at 37°. We inoculated a total of 5µl (1×10^6^ conidia/larvae) to each larva (n=10). We used as the control a group composed of larvae inoculated with 5µl of PBS. We performed inoculations via the last left proleg using a Hamilton syringe (7000.5KH). After infection, we maintained the larvae in petri dishes at 37° C in the dark and scored them daily during ten days. We considered larvae that did not move in response to touch as dead. To evaluate whether there was significant heterogeneity in the virulence profiles of the different strains within each species, we performed log-rank (Mantel-Cox) and Gehan-Breslow-Wilcoxon tests in Graph Pad (Supplementary Table 3).

To measure the antifungal susceptibility of the clinical strains, we applied the EUCAST (European Committee for Antimicrobial Susceptibility Testing) reference method version 9.3.1 (Arendrup et al., 2017). For all strains, we tested their susceptibility to four antifungal drug classes: a) Polyenes: Amphotericin B (AMB, Sigma-Aldrich Quimica, Madrid, Spain); b) Azoles: itraconazole (ICZ, Janssen Pharmaceutica, Madrid, Spain), voriconazole (VCZ, Pfizer SA, Madrid, Spain), posaconazole (PCZ, Schering-Plough Research Institute, Kenilworth, NJ, USA); c) Echinocandins: caspofungin (CPF, Merck & Co. Inc, Rahway, NJ, USA), micafungin (MCF, Astellas Pharma Inc., Tokyo Japan), anidulafungin (AND, Pfizer SA. Madrid, Spain); and d) Allylamine: Terbinafine (TRB, Novartis, Basel, Switzerland). The final concentrations tested ranged from 0.03 to 16 mg/liter for amphotericin B, terbinafine, and caspofungin; from 0.015 to 8 mg/liter for itraconazole, voriconazole and posaconazole; from 0.007 to 4 mg/liter for anidulafungin; and from 0.004 to 2 mg/liter for micafungin. *A. flavus* ATCC 204304 and *A. fumigatus* ATCC 204305 were used as quality control strains in all tests performed. MICs for amphotericin B, itraconazole, voriconazole, posaconazole, and terbinafine, and minimal effective concentrations (MECs) for anidulafungin, caspofungin and micafungin were visually read after 24 and 48 h of incubation at 35°C in a humid atmosphere. To assess the relationship between antifungal susceptibility and strain/species identification, we carried out Principal Component Analysis (PCA) with scaled MIC/MEC values with the R package FactoMineR (Lê et al., 2008), and data visualization with the factoextra v.1.0.6 package.

### Genome sequencing

To understand the genomic similarities and differences within and between these pathogenic *Aspergillus* species and how they are associated with differences in drug susceptibility and virulence profiles, we sequenced the genomes of all 15 strains. Each strain was grown in glucose-yeast extract-peptone (GYEP) liquid medium (0.3% yeast extract and 1% peptone; Difco, Soria Melguizo) with 2% glucose (Sigma-Aldrich, Spain) for 24 h to 48 h at 30°C. After mechanical disruption of the mycelium by vortex mixing with glass beads, genomic DNA of isolates was extracted using the phenol-chloroform method (Holden DW, 1994). The preparation of DNA libraries was performed using the Nextera® TM DNA Library PrepKit (Illumina Inc., San Diego, CA, USA) according to manufacturer’s guidelines. DNA quantification was carried out using the QuantiFluor® dsDNA System and the QuantiFluor® ST Fluorometer (Promega, Madison, WI, USA) and its quality was checked with the Agilent 2100 Bioanalyzer (Agilent Technologies Inc., Santa Clara, CA, USA). Sequencing was performed in the Illumina platform NextSeq500, following the manufacturer’s protocols (Illumina Inc, San Diego, CA, USA). We performed an initial quality analysis of the sequence reads using FastQC, v.0.11.7 (https://www.bioinformatics.babraham.ac.uk/projects/fastqc/). We inspected sequence reads for contaminants using BLAST (Altschul et al., 1990) and MEGAN5 (Huson and Weber, 2013). We trimmed low quality bases (LEADING = 3; TRAILING = 3; SLIDINGWINDOW: windowSize = 4 and requiredQuality = 15), removing both short sequences (< 90 bp) and Nextera adaptors, with Trimmomatic v.0.38 (Bolger et al., 2014).

### Genome assembly and annotation

We assembled the genomes of all strains with SPAdes v3.12.0 (Bankevich et al., 2012). We corrected bases, fixed mis-assemblies, and filled gaps with Pilon, v.1.22 (Walker et al., 2014). We assessed assembly quality using QUAST, v.4.6.3 (Gurevich et al., 2013). We assessed genome assembly completeness using Benchmarking Universal Single-Copy Orthologs (BUSCO) (Simão et al., 2015) and the 4,046 Eurotiomycetes BUSCO gene set (genes from OrthoDB that are thought to be universally single copy). We carried out gene prediction with AUGUSTUS v.3.3.1 (Stanke et al., 2004) using the gene models of *A. fumigatus* as reference. We carried out functional annotation with InterProScan 5.34-73.0 (Jones et al., 2014).

### Orthogroup identification

To identify orthologs (and closely related paralogs) across strains, we performed all-vs-all searches with blastp 2.7.1+ (Altschul et al., 1990) using the strains’ predicted proteomes. We used OrthoFinder v.2.3.3 (Emms and Kelly, 2015) to generate orthogroups using pre-computed BLAST results (-og option) and a Markov Clustering (MCL) inflation value of 1.5. We considered an orthogroup “species-specific” if it possessed one or more protein sequences from only one species.

### Identification of single nucleotide polymorphisms and insertions / deletions

To characterize genetic variation within and between the three pathogenic *Aspergillus* species, we assessed single nucleotide polymorphisms (SNPs) and insertions / deletions (indels). We used BWA-MEM v.0.7.17 (Li and Durbin, 2009) with default parameters to map reads to the reference genome sequences for *A. fumigatus, A. lentulus* and *A. fumigatiaffinis* (CNM-CM8686, CNM-CM7927, and CNM-CM6805, respectively). We did not use type strains as reference genomes for the species under study, because they are not from Spain. Duplicate reads were identified using PICARD MarkDuplicates v.2.9.2 (http://broadinstitute.github.io/picard). We indexed genomes using SAMTOOLS v.1.8 (Li et al., 2009) for subsequent variant detection analyses. We used GenomeAnalysisTK (GATK) v.3.6 for SNP calling with the recommended hard filtering parameters (McKenna et al., 2010; Depristo et al., 2011). We used SnpEff v.4.3t (Cingolani P, Platts A, Wang le L, Coon M, Nguyen T, Wang L, Land SJ, Lu X et al., 2013) to annotate and predict the functional effect of SNPs and indels. We aligned protein and coding sequences for genes of interest with MAFFT v.7.397 (Katoh and Standley, 2013), using the –auto mode. We used Jalview v.2.10.3 (Waterhouse et al., 2009) to visualize SNPs, and a Python script to recover non-synonymous mutations compared to the reference, *A. fumigatus* A1163. Enrichment analysis of GO terms in genes with high impact SNPs and indels for each species was carried out with GOATOOLS v.0.9.9 (Klopfenstein et al., 2018).

### Genetic determinants important for virulence

To examine whether SNPs, indels, and number of paralogs in a given orthogroup were associated with virulence, we recovered 215 genes in *A. fumigatus* Af293 considered genetic determinants of virulence based on their presence in PHI-base (Winnenburg, 2006) and in previously published studies (Abad et al., 2010; Kjærbølling et al., 2018). We obtained functional annotation of these virulence-related genes from FungiDB (Basenko et al., 2018).

### Maximum-likelihood phylogenomics

To reconstruct the evolutionary history of our 15 strains and closely related *Aspergillus* species, we first downloaded or assembled genomes of other strains of the three pathogenic species or their closely relatives that are publicly available. Specifically, we downloaded the genomes of *Aspergillus novofumigatus* IBT16806 (Kjærbølling et al., 2018), *Aspergillus lentulus* IFM 54703^T^ (Kusuya et al., 2016), *Aspergillus fischeri* NRRL181 (Fedorova et al., 2008), *Aspergillus udagawae* IFM46973 (Kusuya et al., 2015), and *Aspergillus viridinutans* FRR_0576 (GenBank accession: GCA_004368095.1). To ensure our analyses also captured the genetic diversity of *A. fumigatus*, we also included additional *A. fumigatus* genomes that spanned the known diversity of *A. fumigatus* strains (Lind et al., 2017). Specifically, we downloaded the genomes of *A. fumigatus* A1163 (Fedorova et al., 2008) and *A. fumigatus* Af293 (Nierman et al., 2005). Additionally, we obtained the raw reads of *A. fumigatus* strains 12-750544 and F16311 (SRA accessions: SRR617737 and ERR769500, respectively). To assemble these genomes, we first quality trimmed the sequence reads using Trimmomatic, v0.36 (Bolger et al., 2014) using parameters described elsewhere (leading:10, trailing:10, slidingwindow:4:20, and minlen:50). The resulting quality-trimmed reads were then used for genome assembly using SPAdes, v3.8.1 (Bankevich et al., 2012), using the ‘careful’ parameter and the ‘cov-cutoff’ parameter set to ‘auto.’ Altogether, we analyzed a total of 24 genomes.

To identify single-copy orthologous genes among the 24 genomes, we implemented the BUSCO, v.2.0.1 (Waterhouse et al., 2013; Simão et al., 2015) pipeline. Specifically, we used the BUSCO pipeline to identify single-copy orthologous genes from genomes using the Eurotiomycetes database of 4,046 orthologs from OrthoDB, v9 (Waterhouse et al., 2013). Among the 4,096 orthologs, we identified 3,954 orthologs with at least 18 taxa represented and aligned the protein sequence each ortholog individually using Mafft, v7.294b (Katoh and Standley, 2013), with the same parameters as described elsewhere (Steenwyk et al., 2019). We then forced nucleotide sequences onto the protein alignment with a custom Python, v3.5.2 (www.python.org), script using BioPython, v1.7 (Cock et al., 2009). The resulting nucleotide alignments were trimmed using trimAl, v1.4 (Capella-Gutierrez et al., 2009), with the ‘gappyout’ parameter. The trimmed alignments were then concatenated into a single matrix with 7,147,728 sites. We then used the concatenated data matrix as input into IQ-TREE, v1.6.11 (Nguyen et al., 2015), with the ‘nbest’ parameter set to 10. The best-fitting model of substitutions was automatically determined using the Bayesian information criterion. The best-fitting model was a general time general time-reversible model with empirical base frequencies, a discrete Gamma model with 4 rate categories, and a proportion of invariable sites (GTR+I+F+G4) (Tavaré, 1986; Yang, 1994; Yang, 1996; Vinet and Zhedanov, 2011). Lastly, we evaluated bipartition support using 5,000 ultrafast bootstrap approximations (Hoang et al., 2018).

In order to build the phylogeny with Cyp51 paralogs, we recovered protein sequences from two orthogroups that included Cyp51A and Cyp51B from *A. fumigatus* Af293 (Afu4g06890 and Afu7g03740, respectively). We generated a maximum-likelihood phylogeny in IQ-Tree v. 1.6.12 (Nguyen et al., 2015), using 1000 Ultrafast Bootstrap Approximation (UFBoot) replicates. The LG+G4 model was chosen as the best according to Bayesian Information Criterion.

### Data availability

All genomes sequenced as part of this work can be accessed through BioProject PRJNA592352; the raw sequence reads are also available through the NCBI Sequence Read Archive. BioSample and Assembly identifiers are presented in Supplementary Table 4. Codes and scripts used in this project are available on the Gitlab repository under https://gitlab.com/SantosRAC/afum_afma_alen2020.

## RESULTS

### Clinical strains show varying antifungal drug susceptibility

By performing PCA on the antifungal drug susceptibility values of all 15 strains, we found that the strains exhibited high heterogeneity in their drug resistance profiles (Figure 1A). In many cases, we found that strains from different species exhibited more similar susceptibility patterns to each other (e.g., strain CNM-CM8686 from *A. fumigatus* with strain CNM-CM6069 from *A. lentulus*) than to other strains from the same species (e.g., strain CNM-CM8686 with strain CNM-CM8057 from *A. fumigatus*), highlighting the magnitude of heterogeneity in drug susceptibility of these species and strains. Principal Component 1 (PC1) explained 37.2% of the variation and separated almost all *A. fumigatus* strains from those of the other two species. Principal Component 2 (PC2) explained 21% of the variation, but did not separate species. The individual contributions of each antifungal drug to each PC are shown in Supplementary Figure 1. We identified that the susceptibility of three antifungals was negatively correlated: micafungin (MCF) *versus* amphotericin B (AMB) and terbinafine (TRB) *versus* AMB (Supplementary Figure 1). In contrast, anidulafungin (AND) and voriconazole (VCZ) were positively correlated (Supplementary Figure 2).

**Figure 1.**
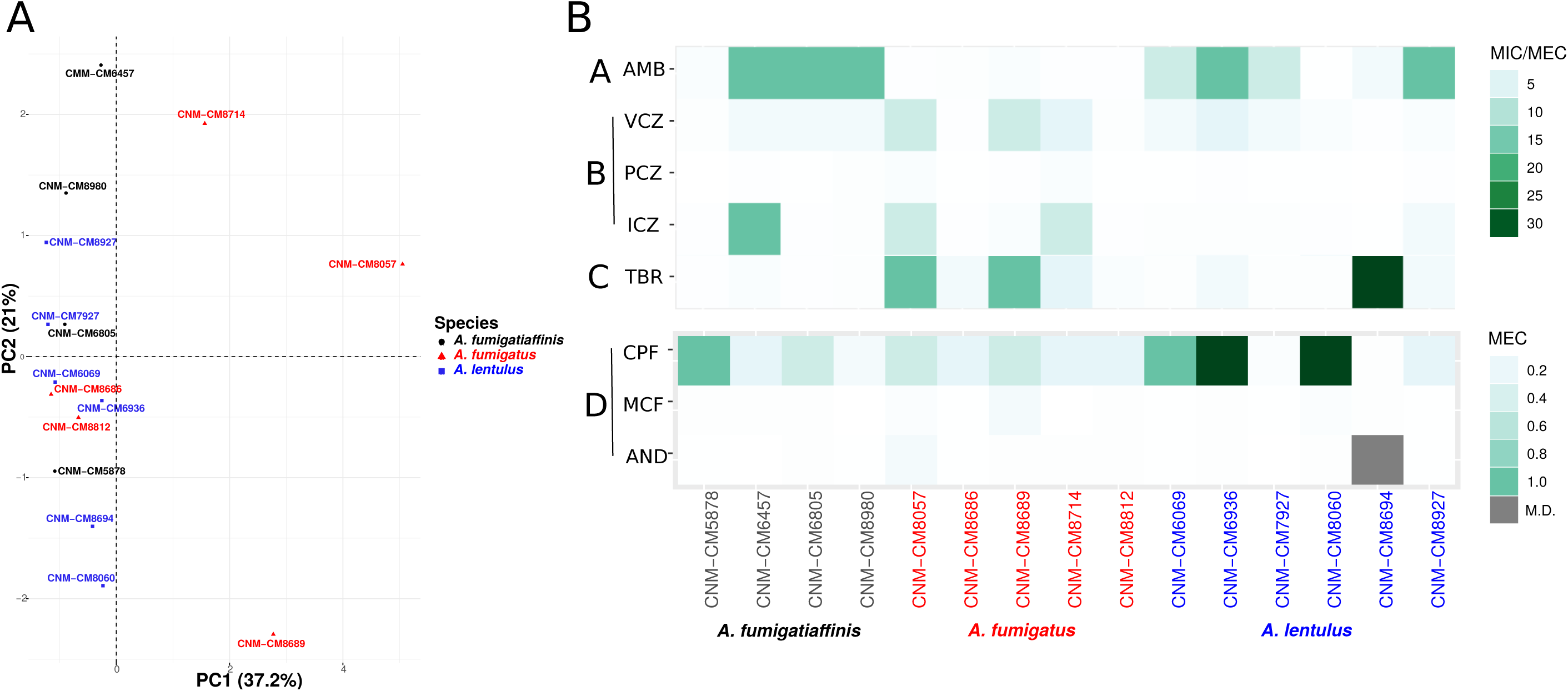
High heterogeneity in drug susceptibility profiles among Spanish strains of three closely related *Aspergillus* pathogens. **(A)** Antifungal susceptibility testing was carried out using EUCAST 9.3. FactoMineR v. 2.1 PCA function was used to scale variables (drug MIC/MECs) and compute Principal Components (PCs), and factoextra v.1.0.6 was used to plot results. PC1 (Dim1) explains most of the variation (37.2% of the variation) and is able to separate *A. fumigatus* from other two species, whereas an overlap is observed in cryptic species (*A. lentulus* and *A. fumigatiaffinis*). **(B)** Antifungal susceptibility testing was carried out using EUCAST 9.3. The Minimum inhibitory concentration (MIC) was obtained for AMB, VCZ, PCZ and ICZ and the Minimum Effective Concentration (MEC) was obtained for TRB, CPF, MCF and AND. A lower scale is shown for echinocandins (bottom panel). Antifungal classes are A: Polyenes; B: Azoles; C: Allylamines; D: Echinocandins. AMB: AmphotericinB; ICZ: itraconazole; VCZ: voriconazole; PCZ: posaconazole; CPF: caspofungin; MCF: micafungin; AND: anidulafungin; TRB: Terbinafine.

We also looked at the differences in susceptibility between strains for each antifungal drug (Figure 1B). Our data show that clinical strains of *A. fumigatus* exhibit lower MICs to AMB compared to *A. lentulus* and *A. fumigatiaffinis*, albeit different levels are observed among different strains (one-way ANOVA; α < 0.05; Tukey multiple comparisons of means for AMB) (Table 1). With the exception of susceptibility of *A. fumigatus* and *A. lentulus* to AMB, for which a significant difference is observed between these two species, we observed high heterogeneity among strains of different species for the other drugs (Table 1). Among azoles, itraconazole (ICZ) and VCZ displayed higher levels of variability across strains. With respect to TRB, the four *A. fumigatiaffinis* strains exhibited low MICs, whereas four *A. fumigatus* strains displayed higher MICs (MIC values >1mg/L) and the other two *A. fumigatus* strains even higher; finally, one *A. lentulus* strain (CNM-CM8694) displayed the highest MICs across all strains (albeit other strains showed in general lower MICs). Among echinocandin drugs, caspofungin (CPF) showed high MECs for the three species. In particular, one strain of *A. fumigatiaffinis* and three of *A. lentulus* were notable in exhibiting very high MECs (MECs ≥ 1mg/L). MECs for MCF and AND were low (≤0.125mg/L) for all strains.

**Table 1.** Susceptibility profile of cryptic *Aspergillus* species isolated in the Mycology Reference Laboratory of Spain.

| Species | Strain Identifier | MIC (mg/L) |  |  |  | MEC (mg/L) |  |  |  |
| --- | --- | --- | --- | --- | --- | --- | --- | --- | --- |
|  |  | AMB | ICZ | VCZ | PCZ | TRB | CPF | MCF | AND |
| *Aspergillus lentulus* | CNM-CM6069 | 8 | 0.5 | 2 | 0.12 | 0.5 | 1 | 0.015 | 0.015 |
|  | CNM-CM6936 | 16 | 0.5 | 4 | 0.25 | 2 | 2 | 0.03 | 0.03 |
|  | CNM-CM7927 | 8 | 0.5 | 2 | 0.12 | 0.5 | 0.06 | 0.015 | 0.007 |
|  | CNM-CM8060 | 0.12 | 0.25 | 1 | 0.12 | 0.5 | 2 | 0.06 | 0.03 |
|  | CNM-CM8694 | 2 | 0.12 | 0.25 | 0.06 | 32 | 0.03 | 0.03 | MD |
|  | CNM-CM8927 | 16 | 2 | 1 | 0.25 | 2 | 0.25 | 0.015 | 0.015 |
| *Aspergillus fumigatiaffinis* | CNM-CM5878 | 1 | 0.25 | 0.5 | 0.06 | 0.25 | 1 | 0.03 | 0.015 |
|  | CNM-CM6457 | 16 | 16 | 2 | 0.25 | 1 | 0.25 | 0.03 | 0.007 |
|  | CNM-CM6805 | 16 | 0.25 | 2 | 0.12 | 0.25 | 0.5 | 0.03 | 0.03 |
|  | CNM-CM8980 | 16 | 0.5 | 2 | 0.5 | 0.5 | 0.12 | 0.007 | 0.015 |
| *Aspergillus fumigatus* | CNM-CM8057 | 0.25 | >8 | >8 | 1 | 16 | 0.5 | 0.06 | 0.12 |
|  | CNM-CM8714 | 0.25 | >8 | 4 | 1 | 4 | 0.25 | 0.007 | 0.03 |
|  | CNM-CM8812 | 0.25 | 0.25 | 0.5 | 0.12 | 1 | 0.25 | 0.03 | 0.03 |
|  | CNM-CM8686 | 0.5 | 0.25 | 0.25 | 0.12 | 2 | 0.25 | 0.015 | 0.015 |
|  | CNM-CM8689 | 1 | 1 | 8 | 0.25 | 16 | 0.5 | 0.125 | 0.03 |
|  | Af293 | 0.5 | 1 | 1 | 0.125 | 2 | 0.125 | 0.007 | 0.007 |
| One-way ANOVA (between species) | P-value | 0.025* | 0.435 | 0.209 | 0.171 | 0.492 | 0.364 | 0.462 | 0.242 |
| Tukey multiple comparisons of means | *Aspergillus fumigatus* - *Aspergillus fumigatiaffinis* | 0.0245507* | - | - | - | - | - | - | - |
|  | *Aspergillus lentulus* - *Aspergillus fumigatiaffinis* | 0.5595621 | - | - | - | - | - | - | - |
|  | *Aspergillus lentulus* - *Aspergillus fumigatus* | 0.0982057 | - | - | - | - | - | - | - |

### Clinical strains show varying levels of virulence

Given functional similarities of the greater wax moth *Galleria mellonella* innate immune system with that of mammals, and prior work showing that moth larvae and mice exhibit similar survival rates when infected with *A. nidulans* (Fernandes et al., 2017), we infected *G. mellonella* larvae with all 15 strains (Figure 2).

**Figure 2.**
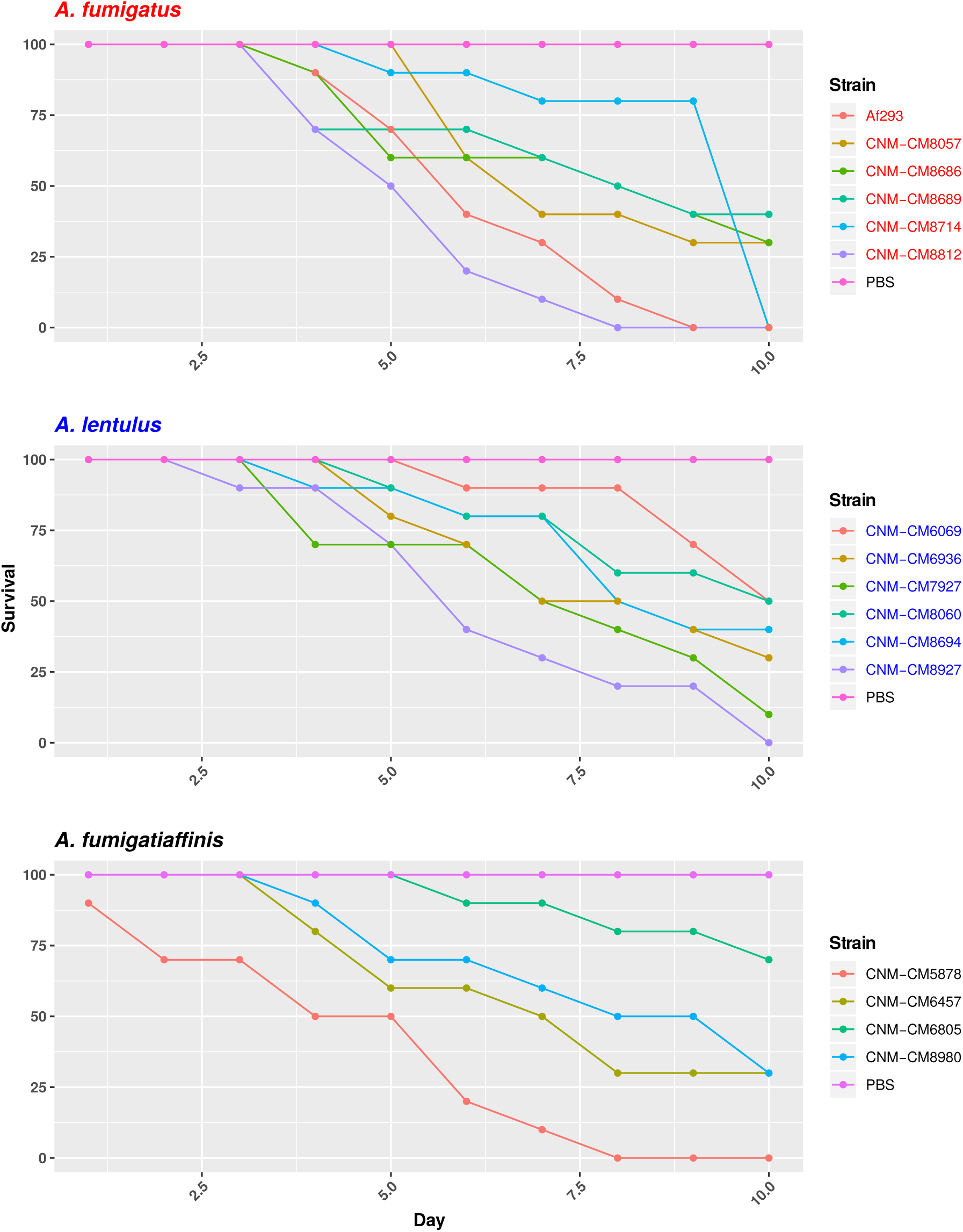
High heterogeneity of virulence levels among Spanish strains of three closely related *Aspergillus* pathogens. Cumulative survival in the model moth *Galleria mellonella*. Larvae infection was carried out via inoculation of 10^6^ conidia. For inoculations, 10 larvae were infected per clinical strain. Experiments were carried out with clinical strains of *A. fumigatus* (upper), *A. lentulus* (middle), and *A. fumigatiaffinis* (lower).

*S*urvival curves revealed high heterogeneity in virulence across clinical strains in the three species (Figure 2). We observed highly virulent strains for which all ten larvae were dead at day 10, such as *A. fumigatus* Af293 (one of our reference strain), *A. fumigatiaffinis* CNM-CM5878 and *A. lentulus* CNM-CM8927. In contrast, other strains were less virulent and >25% larvae survived to the last day of data collection, such as *A. lentulus* CNM-CM6069 and CNM-CM8060. Moreover, we confirmed a high heterogeneity in virulence across strains in the same or in different species (Mantel-Cox and Gehan-Breslow-Wilcoxon tests are shown in the Supplementary Table 3). For example, a similar survival curve is shared between *A. fumigatus* CNM-CM8812 and *A. fumigatiaffinis* CNM-CM5878, whereas *A. fumigatus* CNM-CM8812 and CNM-CM8714 lead to different results.

### Genomic variation within and between Spanish strains of *A. fumigatus, A. lentulus*, and *A. fumigatiaffinis*

To begin exploring the potential genetic underpinnings of species and strain variation in drug susceptibility and virulence, we conducted comparative genomic analyses. The genomes of all 15 strains were of high quality and contained 97-98% of expected complete and single-copy BUSCOs (Supplementary Table 4).

*A. lentulus* and *A. fumigatiaffinis* genomes had larger gene repertoires (9,717 - 9,842 and 10,329 – 10,677, respectively) than *A. fumigatus* (8,837 – 8,938), consistent with previous genome studies of *A. lentulus* and *A. fumigatus* (Nierman et al., 2005; Fedorova et al., 2008; Kusuya et al., 2016). A genome-scale phylogenetic analysis using the nucleotide sequences of BUSCOs with previously sequenced strains (Figure 3A) supports the close relationship between *A. lentulus* and *A. fumigatiaffinis*.

**Figure 3.**
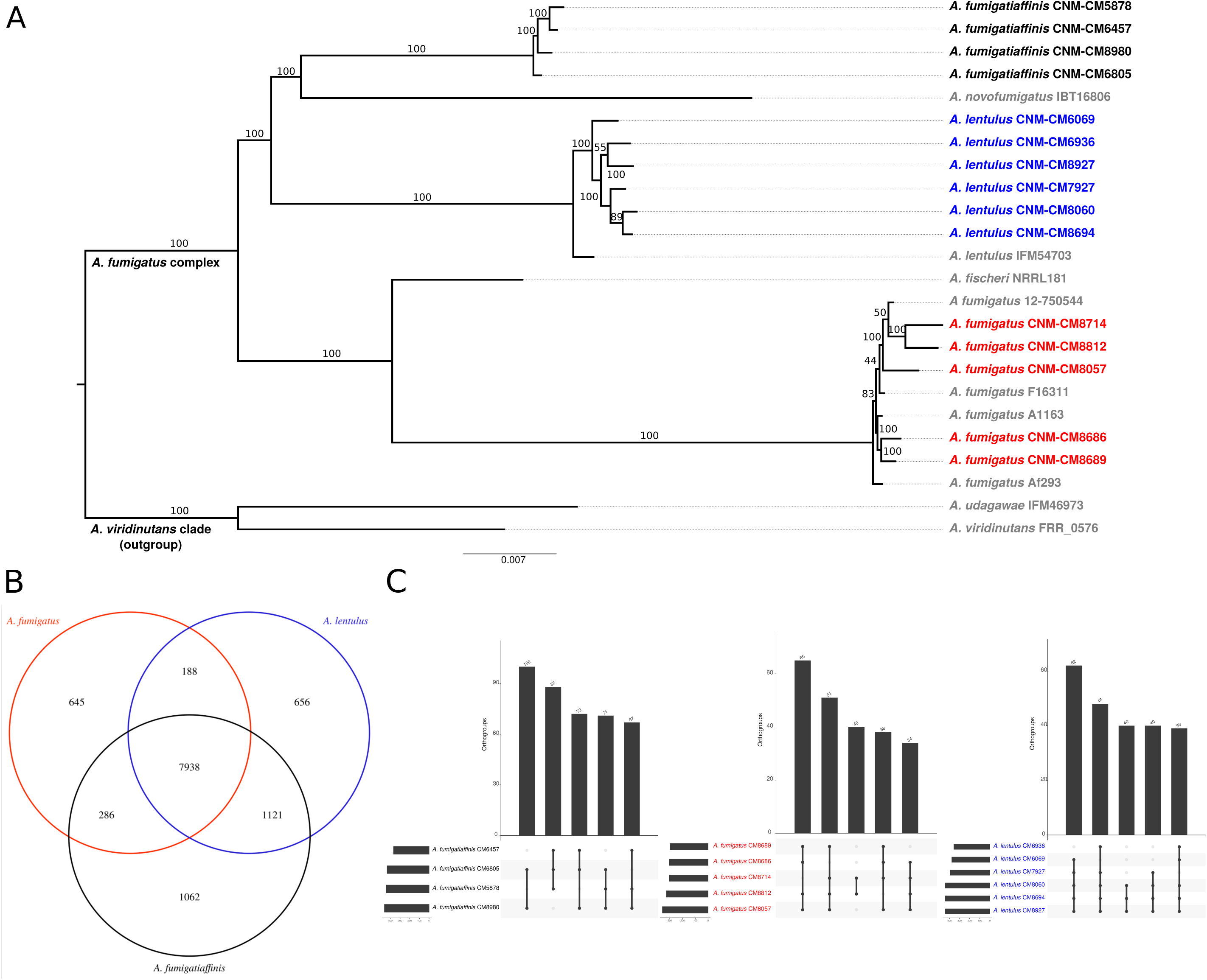
Genomics of the three closely related *Aspergillus* pathogens. **(A)** Genome-scale phylogeny of the section *Fumigati* species used in this study and additional species with sequenced genomes. The *A. viridinutans* clade is presented as a sister clade. Spanish strains sequenced in this work are colored in red (*A. fumigatus*), blue (*A. lentulus*) and black (*A. fumigatiaffinis*). The newly sequenced *A. fumigatiaffinis* strains form a separated group that is closely related to *A. novofumigatus*. All *A. lentulus* strains in this work group together and share an ancestor with *A. lentulus* IFM54703, the only sequenced strain in this species to date. The *A. fumigatus* strains sequenced in this work form different internal groups in the clade with other strains in the species (e.g. strains CNM-CM8714 and CNM-CM8812 group together and strains CNM-CM8686 and CNM-CM8689 form another group). **(B)** *A. fumigatiaffinis* and *A. lentulus* shares the highest number of common orthogroups and *A. fumigatiaffinis* displays the highest number of exclusive orthogroups. OrthoFinder v.2.3.3 was used to group proteins in all clinical strains into orthogroups. *In house* Perl scripts were used to recover orthogroups in each species and VennDiagram in R (Chen and Boutros, 2011) was used to draw intersections. We considered species-specific orthologs those that were present in at least one strain of a given species, with no representative from another species. **(C)** Orthogroups shared by all and “all but one” strains are the most frequent in three closely related *Aspergillus* pathogens. OrthoFinder v.2.3.3 (Emms and Kelly, 2015) was used to group proteins in all clinical strains into orthogroups. *A. lentulus, A. fumigatus* and *A. fumigatiaffinis* presented 9,008, 8,321, and 9,423 orthologous genes present in all strains, respectively. The five largest combination of orthogroups are shown. As expected, the most frequent combination of orthogroups are those in all strains but one. A high number of orthogroups are shared between *A. fumigatus* CNM-CM8714 and CNM-CM8812, reflecting their close phylogenetic relationship.

### Genome diversity among and within species across clinical strains

Examination of orthogroups across the fifteen strains and three species revealed that most genes (7,938) are shared by all three species (Figure 3B). *A. fumigatiaffinis* has a larger set of species-specific genes (1,062) than *A. lentulus* (656) or *A. fumigatus* (645), consistent with its larger genome size and gene number. The numbers of shared genes between *A. lentulus* and *A. fumigatiaffinis* are also higher than intersections between each of them with *A. fumigatus*, consistent with their closer evolutionary affinity (Figure 3A). Within each species, most orthogroups are found in all strains (9,008, 8,321, and 9,423 in *A. lentulus, A. fumigatus*, and *A. fumigatiaffinis*, respectively); approximately 5.4-6.13% of genes in each species appear to vary in their presence between strains (Supplementary Figure 3). Among these, we noted that orthogroups that are present all but one strain are usually the most frequent (Figure 3C).

We identified a total of 114,378, 160,194, and 313,029 SNPs in *A. fumigatus, A. fumigatiaffinis* and *A. lentulus*, respectively. We identified 406, 493, and 747 SNPs in *A. fumigatus, A. fumigatiaffinis* and *A. lentulus*, respectively as high-impact polymorphisms; these polymorphisms are those whose mutation is presumed to be highly deleterious to protein function). Similarly, out of a total of 11,698 (*A. fumigatus*), 20,135 (*A. fumigatiaffinis*) and 34,506 (*A. lentulus*) indels segregating within each species, we identified 615, 1,739, and 1,830 high-impact indels in *A. fumigatus, A. fumigatiaffinis*, and *A. lentulus*, respectively.

Gene Ontology (GO) enrichment analysis was carried out for genes with high impact SNPs and indels (α = 0.05). *A. fumigatus* only showed GO terms identified as underrepresented in “cellular process” and several cellular compartments (“protein-containing complex”, “intracellular organelle part”, “organelle part”, “cytoplasmic part”, “cell part”). *A. lentulus* had “nucleoside metabolic” and “glycosyl compound metabolic processes” enriched, and *A. fumigatiaffinis* showed enriched terms for “modified amino acid binding”, “phosphopantetheine binding”, “amide binding”, “transition metal ion binding”, “zinc ion binding”, “chitin binding”, and “ADP binding”. *A. lentulus* and *A. fumigatiaffinis* genes with high impact SNPs and indels also showed underrepresented GO terms (Supplementary Table 5). We also analysed SNPs and indels separately (Supplementary Table 5).

### Polymorphisms in major antifungal target genes correlate with antifungal susceptibility

Given the observed variation within and between species in antifungal drug susceptibility, we examined DNA sequence polymorphisms in genes known to be involved in antifungal susceptibility to azoles and echinocandins. In particular, we examined patterns of sequence variation in the 14α-sterol demethylase gene *cyp51A* (Afu4g06890) and in the 1,3-beta-glucan synthase catalytic subunit gene *fks1* (Afu6g12400). Using *A. fumigatus* A1163 as reference, we identified important species-and strain-specific polymorphisms in both *cyp51A* and *fks1* (Figure 4A; Table 2 shows a detailed breakdown of all SNP and indel polymorphisms per strain).

**Table 2.** Single-nucleotide polymorphisms and insertions / deletions in *cyp51* family and *fks1* genes in each species individually.

| Species | Polymorphism type | Gene | High impact variant | Low impact variant | Moderate impact variant | Modifier impact variant |
| --- | --- | --- | --- | --- | --- | --- |
| *A. fumigatus* | INDELs | *cyp51A* | 0 | 0 | 0 | 6 |
| *A. fumigatus* | SNPs | *cyp51A* | 0 | 0 | 5 | 23 |
| *A. lentulus* | INDELs | *cyp51A* | 0 | 0 | 0 | 10 |
| *A. lentulus* | SNPs | *cyp51A* | 0 | 8 | 3 | 92 |
| *A. fumigatus* | INDELs | *cyp51B* | 0 | 0 | 0 | 1 |
| *A. fumigatus* | SNPs | *cyp51B* | 0 | 1 | 1 | 7 |
| *A. lentulus* | INDELs | *cyp51B* | 0 | 0 | 0 | 1 |
| *A. lentulus* | SNPs | *cyp51B* | 0 | 0 | 0 | 2 |
| *A. fumigatiaffinis* | INDELs | *cyp51C* | 0 | 0 | 0 | 28 |
| *A. fumigatiaffinis* | SNPs | *cyp51C* | 0 | 10 | 1 | 157 |
| *A. fumigatiaffinis* | INDELs | *fks1* | 0 | 0 | 0 | 45 |
| *A. fumigatiaffinis* | SNPs | *fks1* | 0 | 23 | 5 | 143 |
| *A. fumigatus* | INDELs | *fks1* | 0 | 0 | 0 | 3 |
| *A. fumigatus* | SNPs | *fks1* | 0 | 4 | 0 | 10 |
| *A. lentulus* | INDELs | *fks1* | 4 | 0 | 0 | 50 |
| *A. lentulus* | SNPs | *fks1* | 0 | 43 | 5 | 218 |

**Figure 4.**
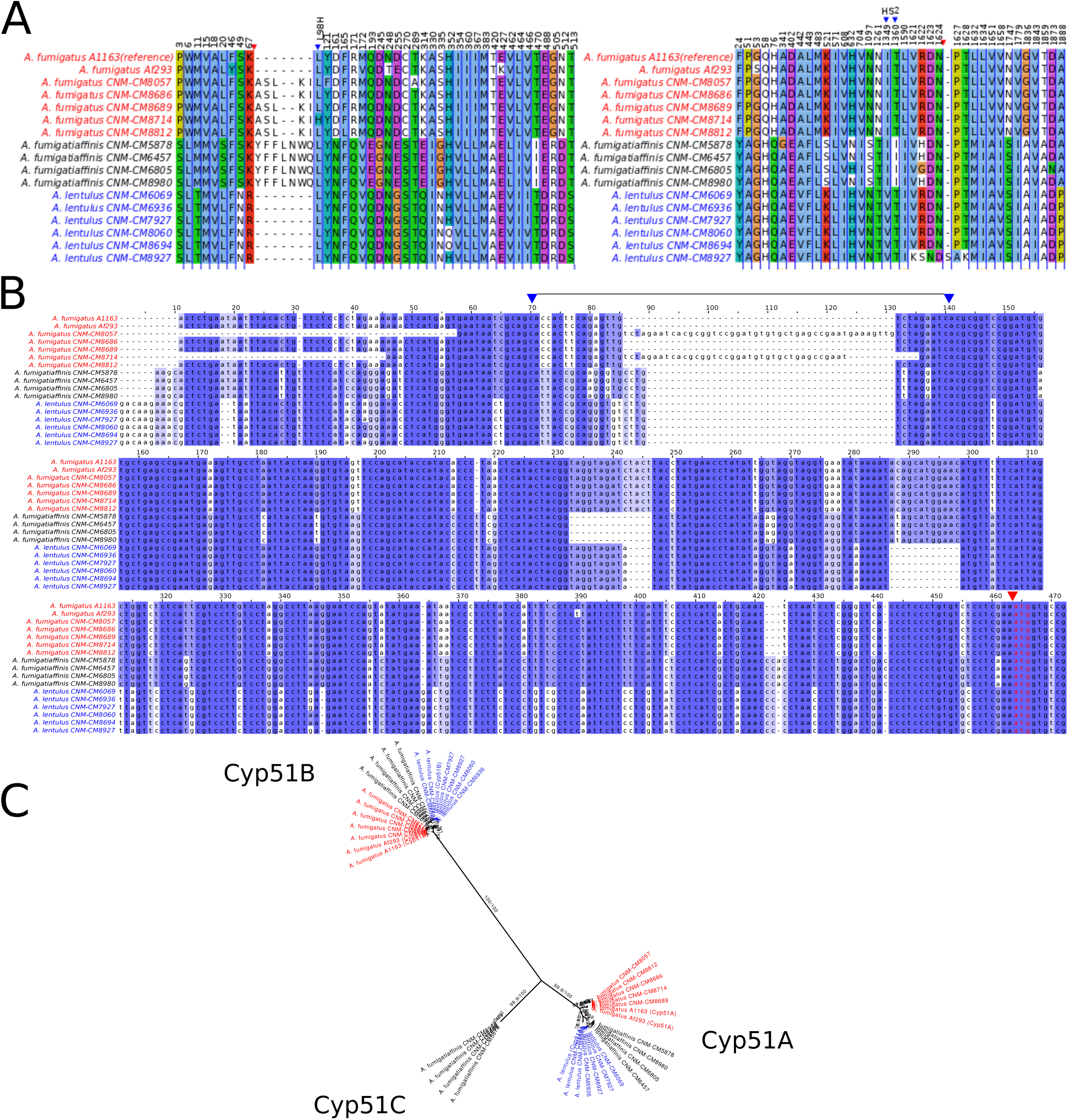
Changes in important genes related to antifungal susceptibility in the three *Aspergillus* pathogens. **(A)** Products of genes related to antifungal resistance, *Cyp51A* (azoles) and *Fks1* (echinocandins), display species- and strain-specific polymorphisms. MAFFT v.7.397 (--auto mode) was used to align protein sequences. Alignments were imported in Jalview v.2.10.3 (Waterhouse et al., 2009), colored in Clustal mode, and non-polymorphic positions were hidden. Only the positions with changes in at least one strain are shown (substitutions or insertions/deletions). Blue triangles highlight amino acid changes in position 98 in Cyp51A and in hot spot 2 (HS2) of *Fks1*. Red triangles indicate insertions/deletions. **(B)** Promoter region of the *cyp51A* gene displays strain-specific mutations among Spanish strains of three closely related *Aspergillus* pathogens. A *in house* python script was used to recover 400 bp upstream to the *cyp51A* gene and the three initial codons. MAFFT v.7.397 (--auto mode) was used to align the nucleotide sequences in all clinical strains. Well-known Tandem Repeat regions in antifungal-resistant strains of *A. fumigatus* are shown between positions 70-140 in the alignment (i.e. TR34 and TR46, observed in CNM-CM8714 and CNM-CM8057, respectively, delimited by two blue arrows in upper part). Polymorphisms in cryptic species were also identified, for instance, the short deletions exclusively found in the cryptic species (either in *A. fumigatiaffinis* or *A. lentulus*) around positions 230-250. Red arrow and red font indicates the start codon. **(C)** Phylogeny of Cyp51 gene family (protein sequences) reveals three different members (Cyp51A, Cyp51B, and the putative Cyp51C) in *A. fumigatiaffinis*. A maximum-likelihood phylogeny was generated in IQ-Tree (v. 1.6.12), using 1000 Ultrafast Bootstrap Approximation (UFBoot) replicates. LG+G4 model was chosen as the best according to Bayesian Information Criterion (BIC). UFBoot and SH-aLRT support values are shown.

An alignment of Cyp51A protein sequences in the three species shows possible insertions in different sites (Figure 4A - red arrow). We observed substitutions in at least one of the clinical strains in the three species in 42 positions that might be correlated with the strains’ varying drug susceptibility levels. For instance, Cyp51A in *A. fumigatus* CNM-CM8714 revealed a well-documented substitution related to azole resistance at position 98 (L98H) (Figure 4A - blue arrows), which might be correlated to its lower susceptibility to ICZ or VCZ compared to other *A. fumigatus* strains (Figure 1B).

We also looked at the promoter region of the *cyp51A* gene (Figure 4B) and identified the Tandem Repeat (TR) insertions TR34 and TR46, previously reported in antifungal resistant strains (Dudakova et al., 2017). These changes were specific to certain clinical strains of *A. fumigatus*. For example, *A. fumigatus* CNM-CM8714 carries the TR34 insertions whereas *A. fumigatus* CNM-CM8057 has a TR46 (region highlighted between blue arrows). There are other variations (short indels) that were exclusive to either *A. lentulus* or *A. fumigatiaffinis*, or both.

Examination of the Fks1 protein sequence alignment from strains of the three species also revealed substitutions in 39 sites (Figure 4A). We also observed an insertion at position 1,626 of *A. lentulus* CNM-CM8927 (red arrow). Fks1 also showed substitutions at positions comprising an important hot-spot 2 (HS2) (blue arrows): all *A. lentulus* strains have a substitution at position 1,349 (I1349V) and all *A. fumigatiaffinis* have a substitution at position 1,360 (T1360I).

Examination of orthogroups revealed that the orthogroup that includes the *cyp51A* gene (Afu4g06890) contained additional paralogs of the *cyp51* family. Thus, we carried out a phylogenetic analysis with the amino acid sequences with the orthogroups containing *cyp51A* and *cyp51B* genes in *A. fumigatus* Af293 (Figure 4C) that comprises the three species in this work. We observed three well-defined clades. The *A. fumigatiaffinis* paralog related to *cyp51A* is likely to represent *cyp51C*, which has been previously reported in other *Aspergillus* species (Hagiwara et al., 2016). Sequence identity between the putative Cyp51C protein in *A. fumigatiaffinis* CNM-CM6805 and Cyp51C (XM_002383890.1) and Cyp51A (XM_002375082.1) of *A. flavus* (Liu et al., 2012) is 471/512 (92%) and 391/508 (77%), respectively.

### Genetic determinants involved in virulence: single-nucleotide polymorphisms, insertions / deletions across strains and within species conservation

To explore the genetic underpinnings of the observed strain heterogeneity in virulence we next examined the SNPs and indels in 215 genes that have previously been characterized as genetic determinants of virulence in *A. fumigatus* (Supplementary Table 6).

Most virulence genetic determinants (146 genes) were found in single-copy in all strains (Supplementary Table 7), whereas 57 genes varied in their number of paralogs across clinical strains (Figure 5). We also identified four virulence determinants that had no orthologs in either *A. lentulus* or *A. fumigatiaffinis*, such as Afu6g07120 (*nudC*), which is an essential protein involved in nuclear movement (Morris et al., 1998), and considered an essential gene in *A. fumigatus* (Hu et al., 2007). Interestingly, we noted 17 virulence determinants that are present in *A. fumigatus* and *A. fumigatiaffinis* but absent in *A. lentulus* (Figure 5 – top panel), such as Afu8g00200 (*ftmD*), one of the genes in the fumitremorgin biosynthetic gene cluster (Abad et al., 2010).

**Figure 5.**
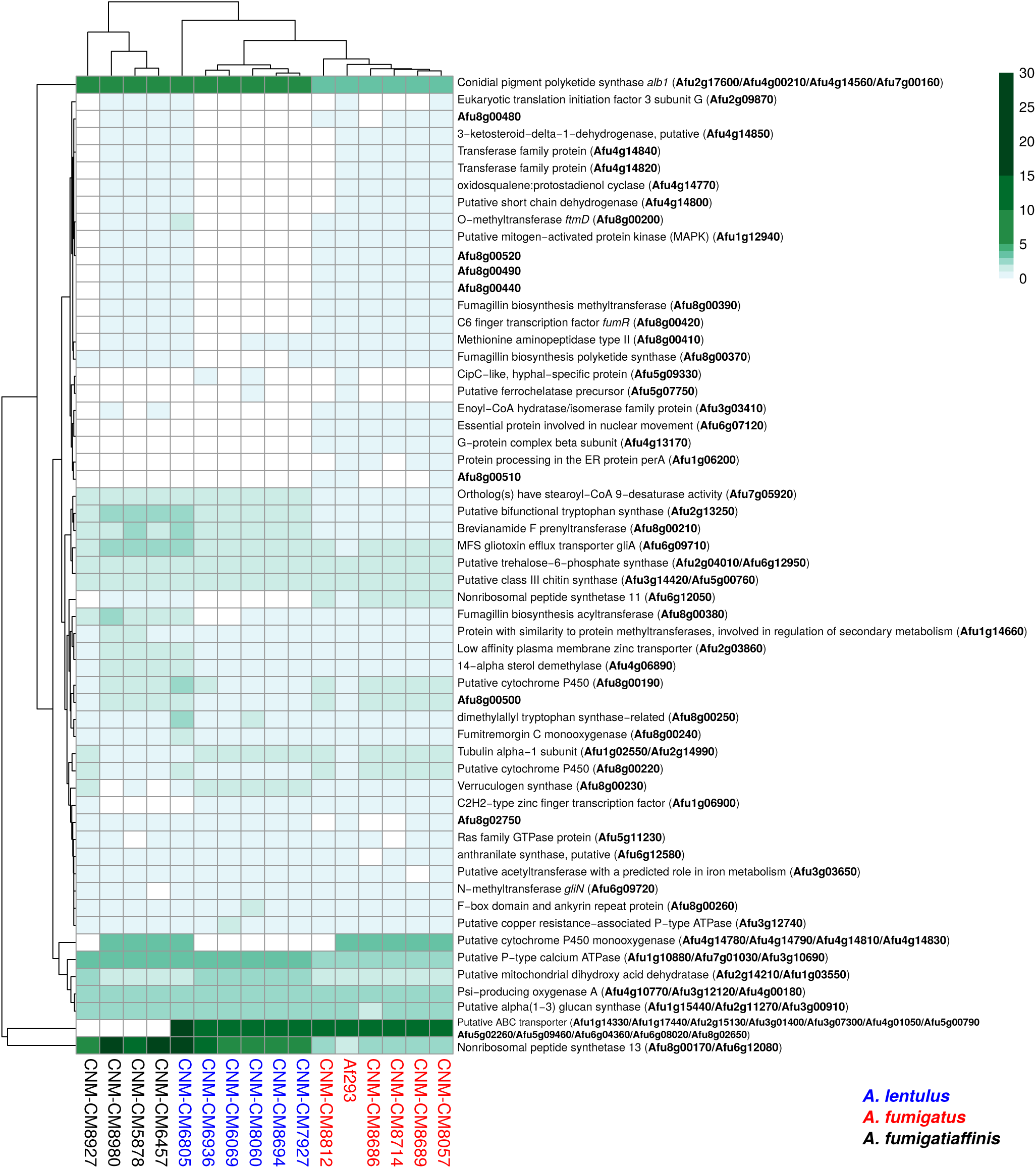
Orthogroups for virulence determinants reveals variable number of paralogs among the three closely related *Aspergillus* pathogens. We searched for 215 known genetic determinants of virulence in *A. fumigatus* Af293 in the species of interest and found they were grouped into 203 orthogroups. 146/203 were found in single copy across all strains and are not shown here. The cladogram above the species reflects similarities between strain presence/absence patterns. *A. fumigatus* Af293 shows a different pattern compared to other strains of *A. fumigatus*, grouping with one of the *A. lentulus* strains (CNM-CM8927). This may reflect the phylogenetic divergence of *A. fumigatus* strain Af293 from other species members. Conidial pigment polyketide synthase *alb1* (Afu2g17600) is one of the genetic determinants of virulence with highest number of copies in cryptic species (n=7) when compared to *A. fumigatus* strains (n=4). Gene identifiers in *A. fumigatus* Af293 are highlighted in bold.

Several virulence determinants exhibited larger numbers of paralogs in one or more species. For example, the conidial pigment polyketide synthase *alb1* (Afu2g17600), which is involved in conidial morphology and virulence (Tsai et al., 1998), is one of the determinants with highest number of paralogs in *A. lentulus* and *A. fumigatiaffinis* (n=7) when compared to *A. fumigatus* strains (n=4). For determinants that contained a gene in at least one strain, we tested correlations between number of paralogs and virulence (lethal time 50: day at which 50% of the larvae were dead, or “ND-end”: the number of dead larvae at the end of the experiment) and we observed no significant correlation suggesting paralog number does not associate with virulence.

## DISCUSSION

*A. fumigatus* and the closely related species *A. lentulus* and *A. fumigatiaffinis* are important causal agents of aspergillosis (Zbinden et al., 2012; Lamoth, 2016). Importantly, the emergence of antifungal resistance is of increasing worldwide concern (Fisher et al., 2018) and antifungal resistant strains of *A. lentulus*, and *A. fumigatiaffinis* (Alastruey-Izquierdo et al., 2014) have been identified. Heterogeneity in virulence across different strains of *A. fumigatus* has also been known for some time (Mondon et al., 1996). Analyses of strain phenotypic and genetic heterogeneity allow us to identify correlations between phenotype and genotype in strains of *Aspergillus* pathogens.

We found that high heterogeneity exists in drug susceptibility and virulence across different strains of *A. fumigatus, A. lentulus*, and *A. fumigatiaffinis*. For one specific antifungal drug, amphotericin B, our results confirmed previous findings that *A. fumigatus* is more susceptible to AMB than strains of cryptic species (Balajee et al., 2004). Studies on the intrinsic resistance to AMB reported for *A. terreus* highlight the importance of stress response pathways, in particular heat shock proteins (such as Hsp90 and Hsp70), as well as enzymes detoxifying reactive oxygen species (Posch et al., 2018). Future work involving genomics on the cryptic species will be able to exploit changes in genetic determinants involved in AMB susceptibility, although this drug is not commonly used in clinical settings.

Regarding virulence, previous work (Sugui et al., 2014) compared survival curves between different species of *Aspergillus* and concluded that *A. fumigatus* is significantly more virulent than *A. lentulus* based on type strains. However, using the *Galleria* model of fungal infection we found that observed within-species variance is greater than it would be intuitively expected for different species. While additional testing using diverse models of fungal disease will be required to test the validity of these observations, our findings reinforce the emerging view that examining within-species variation in *Aspergillus* pathogens is an important, yet poorly studied and understood, dimension of fungal virulence.

The advent of whole genome sequencing boosted our understanding of the biology of the genus *Aspergillus* (de Vries et al., 2017). Several works have previously analysed genomic data of *A. fumigatus* strains (Abdolrasouli et al., 2015; Takahashi-Nakaguchi et al., 2015), uncovering *cyp51A* mutations in *A. fumigatus* populations (Abdolrasouli et al., 2015). Some studies have also focused on antifungal drug susceptibility (Garcia-Rubio et al., 2018) or virulence potential (Puértolas-Balint et al., 2019) using only the genome sequencing data for strains of *A. fumigatus*. Correlations between phenotypic traits, such as antifungal susceptibility or virulence and genetic traits have been also studied in other well-studied pathogens, such as the opportunistic yeast *Candida albicans* (Hirakawa et al., 2015). However, to our knowledge, this is the first study that examines the phenotypic and genetic heterogeneity among strains of species closely related to *A. fumigatus*.

The three main classes of antifungal drugs comprise polyenes, azoles, and echinocandins, involved in ergosterol composition of fungal membrane, ergosterol biosynthesis, and the cell wall biopolymer (1,3)-β-D-glucan, respectively (Robbins et al., 2017). Due to toxicity to host cells, polyenes are only used in exceptional cases, and first-line prophylaxis and treatment of aspergillosis is usually carried out with azoles (Garcia-Rubio et al., 2017; Garcia-Vidal et al., 2019). In *A. fumigatus*, research has focused on azole susceptibility testing and correlation with point mutations in the *cyp51A* gene and tandem repeat insertions in its promoter region (Chen et al., 2020; Zakaria et al., 2020). Studies on echinocandins have focused on the (1,3)-β-D-glucan synthase enzyme, encoded by the *fks1* gene (Robbins et al., 2017). Particularly, two hot-spots have been studied (Gonçalves et al., 2016). Although most studies report mutations in *Candida*, previous work reported point mutations in *fks1* hot spot 1 associated with echinocandin resistance in *A. fumigatus* (Jiménez-Ortigosa et al., 2017). This work did not find changes among *A. fumigatus* clinical strains, but we did observe changes in hot spot 2 that were specific to the cryptic species; further examination of these changes with respect to echinocandin susceptibility is an interesting future avenue of research.

Major changes in protein Cyp51A that correlated with azole resistance include the TR34/L98H and the TR46/Y121F/T289A changes (Beardsley et al., 2018). In previous studies, alterations such as the insertion of TR34 and TR46 (Dudakova et al., 2017) were only found in the *cyp51A* promoter of *A. fumigatus* strains. However, as Dudakova et al. previously mentioned, “data on *cyp51* promoter alterations are scarce” outside *A. fumigatus* (Dudakova et al., 2017). Our work explored the promoter region of the *cyp51A* gene in *A. lentulus* and *A. fumigatiaffinis*, two closely related pathogenic species, and identified these promoter region changes only in two strains of *A. fumigatus* and not in either of the two cryptic species. We also identified other changes in proteins encoded by *cyp51A* and *fks1* that can be used in the future to generate mutants and test the effect of mutations in well-studied wild-type strains of *A. fumigatus* (Chen et al., 2020).

The evolution of the gene families that contain genes involved in drug resistance might also give us clues on how drug resistance evolves in fungal populations; previous studies (Hawkins et al., 2014; Zheng et al., 2019) report two paralogs of c*yp51* in *Aspergillus fumigatus, Penicillium digitatum, Magnaporthe oryzae* and *A. nidulans*, while *Fusarium graminearum* and *A. flavus* have three *cyp51* genes (Dudakova et al., 2017). Interestingly, our study found a paralog of the *cyp51A* gene in *A. fumigatiaffinis* that likely corresponds to *cyp51C*. Whole-genome sequence analysis in *A. flavus* reported substitutions in the three paralogous genes (*cyp51A, cyp51B*, and *cyp51C*) in the context of antifungal resistance (Sharma et al., 2018). Novel substitutions identified in *cyp51C* and modelling of protein changes suggested possible effects on drug binding. Next steps in studies of azole susceptibility in *A. fumigatiaffinis* strains could include the analysis of this putative *cyp51C* gene and its role in the organism’s observed drug susceptibility profile.

While we extended the understanding of strain heterogeneity to strains in the three different species of *Aspergillus* pathogens in terms of virulence and drug susceptibility, phylogenomic analyses support the between-species groupings (e.g. closer relationship between *A. lentulus* and *A. fumigatiaffinis* than each of these species to *A. fumigatus*) inferred when using single phylogenetic markers (e.g., from analyses of *benA*). While the analysis of disease-relevant genes (e.g., *cyp51A* and fks*1*) and patterns of paralogous gene number of the genetic determinants of virulence support the relationships depicted in phylogenies, we see many strain-specific changes. Our work is a first opportunity to exploit these specific variants that must be related to variance in virulence and drug susceptibility (if the source is genetic variation).

Albeit this study focused on substitutions in genes *cyp51A* and *fks1*, there is also increasing research on non-*cyp51* (Zakaria et al., 2020) and non-*fks1* (Szalewski et al., 2018) genetic changes. Future exploitation of genomic data on strains of *A. fumigatus* and closely related species could also exploit these additional genes. These future studies could also exploit new antifungal drugs, such as olorofim, which has also been tested on cryptic species of *Aspergillus* (Rivero-Menendez et al., 2019). Finally, future phenotypic and genomic analyses can help us to better understand more complex topics involving antifungal drugs, such as resistance, persistence and tolerance (as well as the role of tolerance in resistance) (Berman and Krysan, 2020). Given possible emergence of antifungal resistance in agriculture (Hawkins et al., 2019), future work could also exploit correlations in antifungals and the origin of these isolates.

## AUTHOR CONTRIBUTIONS

R.A.C.S., J.L.S., M.E.M., A.A.I., G.H.G., and A.R. designed the experiments. L.P., O.R.M. and R.W.B. performed experiments. R.A.C.S. and J.L.S. ran bioinformatic analyses. R.A.C.S., M.E.M., J.L.S. and A.R. wrote the manuscript. All authors revised the manuscript.

## ACKNOWLEDGEMENTS

This work used resources of the “Centro Nacional de Processamento de Alto Desempenho em São Paulo (CENAPAD-SP). Computational infrastructure was provided by The Advanced Computing Center for Research and Education (ACCRE) at Vanderbilt University.

## REFERENCES

Abad, A., Victoria Fernández-Molina, J., Bikandi, J., Ramírez, A., Margareto, J., Sendino, J., et al. (2010). What makes Aspergillus fumigatus a successful pathogen? Genes and molecules involved in invasive aspergillosis. Rev. Iberoam. Micol. doi:10.1016/j.riam.2010.10.003.

Abdolrasouli, A., Rhodes, J., Beale, M. A., Hagen, F., Rogers, T. R., Chowdhary, A., et al. (2015). Genomic context of azole resistance mutations in Aspergillus fumigatus determined using whole-genome sequencing. MBio. doi:10.1128/mBio.00536-15.

Agarwal, R., Chakrabarti, A., Shah, A., Gupta, D., Meis, J. F., Guleria, R., et al. (2013). Allergic bronchopulmonary aspergillosis: Review of literature and proposal of new diagnostic and classification criteria. Clin. Exp. Allergy. doi:10.1111/cea.12141.

Alastruey-Izquierdo, A., Alcazar-Fuoli, L., and Cuenca-Estrella, M. (2014). Antifungal susceptibility profile of cryptic species of aspergillus. Mycopathologia. doi:10.1007/s11046-014-9775-z.

Altschul, S. F., Gish, W., Miller, W., Myers, E. W., and Lipman, D. J. (1990). Basic local alignment search tool. J. Mol. Biol. doi:10.1016/S0022-2836(05)80360-2.

Arendrup MC, Meletiadis J, Mouton JW, Lagrou K, Howard SJ, Subcommittee on Antifungal Susceptibility Testing of the ESCMID European Committee for Antimicrobial Susceptibility Testing. 2017. EUCAST DEFINITIVE DOCUMENT E.DEF 9.3.1: Method for the determination of broth dilution minimum inhibitory concentrations of antifungal agents for conidia forming moulds. http://www.eucast.org/fileadmin/src/media/PDFs/EUCAST_files/AFST/Files/EUCAST_E_Def_9_3_1_Mould_testing__definitive.pdf

Balajee, S. A., Gribskov, J. L., Hanley, E., Nickle, D., and Marr, K. A. (2005). Aspergillus lentulus sp. nov., a new sibling species of A. fumigatus. Eukaryot. Cell. doi:10.1128/EC.4.3.625-632.2005.

Balajee, S. A., Kano, R., Baddley, J. W., Moser, S. A., Marr, K. A., Alexander, B. D., et al. (2009). Molecular identification of Aspergillus species collected for the transplant-associated infection surveillance network. J. Clin. Microbiol. doi:10.1128/JCM.01070-09.

Balajee, S. A., Weaver, M., Imhof, A., Gribskov, J., and Marr, K. A. (2004). Aspergillus fumigatus Variant with Decreased Susceptibility to Multiple Antifungals. Antimicrob. Agents Chemother. doi:10.1128/AAC.48.4.1197-1203.2004.

Bankevich, A., Nurk, S., Antipov, D., Gurevich, A. A., Dvorkin, M., Kulikov, A. S., et al. (2012). SPAdes: A new genome assembly algorithm and its applications to single-cell sequencing. J. Comput. Biol. doi:10.1089/cmb.2012.0021.

Basenko, E. Y., Pulman, J. A., Shanmugasundram, A., Harb, O. S., Crouch, K., Starns, D., et al. (2018). FungiDB: An integrated bioinformatic resource for fungi and oomycetes. J. Fungi. doi:10.3390/jof4010039.

Beardsley, J., Halliday, C. L., Chen, S. C. A., and Sorrell, T. C. (2018). Responding to the emergence of antifungal drug resistance: Perspectives from the bench and the bedside. Future Microbiol. doi:10.2217/fmb-2018-0059.

Berman, J., and Krysan, D. J. (2020). Drug resistance and tolerance in fungi. Nat. Rev. Microbiol. doi:10.1038/s41579-019-0322-2.

Bolger, A. M., Lohse, M., and Usadel, B. (2014). Trimmomatic: A flexible trimmer for Illumina sequence data. Bioinformatics. doi:10.1093/bioinformatics/btu170.

Brown, G. D., Denning, D. W., Gow, N. A. R., Levitz, S. M., Netea, M. G., and White, T. C. (2012). Hidden killers: Human fungal infections. Sci. Transl. Med. doi:10.1126/scitranslmed.3004404.

Capella-Gutierrez, S., Silla-Martinez, J. M. & Gabaldon, T. trimAl: a tool for automated alignment trimming in large-scale phylogenetic analyses. Bioinformatics 25, 1972–1973 (2009).

Chen, P., Liu, M., Zeng, Q., Zhang, Z., Liu, W., Sang, H., et al. (2020). Uncovering New Mutations Conferring Azole Resistance in the Aspergillus fumigatus cyp51A Gene. Front. Microbiol. doi:10.3389/fmicb.2019.03127.

Cingolani P, Platts A, Wang le L, Coon M, Nguyen T, Wang L, Land SJ, Lu X, R. D., Cingolani, P., Platts, A., Wang, L. L. L., Coon, M., Nguyen, T., et al. (2013). A program for annotating and predicting the effects of single nucleotide polymorphisms, SnpEff: SNPs in the genome of Drosophila melanogaster strain w1118; iso-2; iso-3. Fly (Austin). doi:10.1016/S1877-1203(13)70353-7.

Cock, P. J. A. et al. Biopython: freely available Python tools for computational molecular biology and bioinformatics. Bioinformatics 25, 1422–1423 (2009)

de Vries, R. P., Riley, R., Wiebenga, A., Aguilar-Osorio, G., Amillis, S., Uchima, C. A., et al. (2017). Comparative genomics reveals high biological diversity and specific adaptations in the industrially and medically important fungal genus Aspergillus. Genome Biol. doi:10.1186/s13059-017-1151-0.

Denning, D. W., Cadranel, J., Beigelman-Aubry, C., Ader, F., Chakrabarti, A., Blot, S., et al. (2016). Chronic pulmonary aspergillosis: Rationale and clinical guidelines for diagnosis and management. Eur. Respir. J. doi:10.1183/13993003.00583-2015.

Depristo, M. A., Banks, E., Poplin, R., Garimella, K. V., Maguire, J. R., Hartl, C., et al. (2011). A framework for variation discovery and genotyping using next-generation DNA sequencing data. Nat. Genet. doi:10.1038/ng.806.

Dudakova, A., Spiess, B., Tangwattanachuleeporn, M., Sasse, C., Buchheidt, D., Weig, M., et al. (2017). Molecular tools for the detection and deduction of azole antifungal drug resistance phenotypes in Aspergillus species. Clin. Microbiol. Rev. doi:10.1128/CMR.00095-16.

Emms, D. M., and Kelly, S. (2015). OrthoFinder: solving fundamental biases in whole genome comparisons dramatically improves orthogroup inference accuracy. Genome Biol. doi:10.1186/s13059-015-0721-2.

Fedorova, N. D., Khaldi, N., Joardar, V. S., Maiti, R., Amedeo, P., Anderson, M. J., et al. (2008). Genomic islands in the pathogenic filamentous fungus Aspergillus fumigatus. PLoS Genet. doi:10.1371/journal.pgen.1000046.

Fernandes, C., Fonseca, F., Goldman, G., Pereira, M., and Kurtenbach, E. (2017). A Reliable Assay to Evaluate the Virulence of Aspergillus nidulans Using the Alternative Animal Model Galleria mellonella (Lepidoptera). BIO-PROTOCOL. doi:10.21769/bioprotoc.2329.

Fisher, M. C., Hawkins, N. J., Sanglard, D., and Gurr, S. J. (2018). Worldwide emergence of resistance to antifungal drugs challenges human health and food security. Science (80-.). doi:10.1126/science.aap7999.

Fuchs, B. B., O’Brien, E., Khoury, J. B. E., and Mylonakis, E. (2010). Methods for using Galleria mellonella as a model host to study fungal pathogenesis. Virulence. doi:10.4161/viru.1.6.12985.

Fuller, K. K., Cramer, R. A., Zegans, M. E., Dunlap, J. C., and Loros, J. J. (2016). Aspergillus fumigatus photobiology illuminates the marked heterogeneity between isolates. MBio. doi:10.1128/mBio.01517-16.

Garcia-Rubio, R., Alcazar-Fuoli, L., Monteiro, M. C., Monzon, S., Cuesta, I., Pelaez, T., et al. (2018). Insight into the significance of aspergillus fumigatus cyp51A polymorphisms. Antimicrob. Agents Chemother. doi:10.1128/AAC.00241-18.

Garcia-Rubio, R., Cuenca-Estrella, M., and Mellado, E. (2017). Triazole Resistance in Aspergillus Species: An Emerging Problem. Drugs. doi:10.1007/s40265-017-0714-4.

Garcia-Vidal, C., Alastruey-Izquierdo, A., Aguilar-Guisado, M., Carratalà, J., Castro, C., Fernández-Ruiz, M., et al. (2019). Executive summary of clinical practice guideline for the management of invasive diseases caused by Aspergillus: 2018 Update by the GEMICOMED-SEIMC/REIPI. Enferm. Infecc. Microbiol. Clin. doi:10.1016/j.eimc.2018.03.018.

Gonçalves, S. S., Souza, A. C. R., Chowdhary, A., Meis, J. F., and Colombo, A. L. (2016). Epidemiology and molecular mechanisms of antifungal resistance in Candida and Aspergillus. Mycoses. doi:10.1111/myc.12469.

Gurevich, A., Saveliev, V., Vyahhi, N., and Tesler, G. (2013). QUAST: Quality assessment tool for genome assemblies. Bioinformatics. doi:10.1093/bioinformatics/btt086.

Hagiwara, D., Watanabe, A., Kamei, K., and Goldman, G. H. (2016). Epidemiological and genomic landscape of azole resistance mechanisms in Aspergillus fungi. Front. Microbiol. doi:10.3389/fmicb.2016.01382.

Hawkins, N. J., Bass, C., Dixon, A., and Neve, P. (2019). The evolutionary origins of pesticide resistance. Biol. Rev. doi:10.1111/brv.12440.

Hawkins, N. J., Cools, H. J., Sierotzki, H., Shaw, M. W., Knogge, W., Kelly, S. L., et al. (2014). Paralog re-emergence: A novel, historically contingent mechanism in the evolution of antimicrobial resistance. Mol. Biol. Evol. doi:10.1093/molbev/msu134.

Hirakawa, M. P., Martinez, D. A., Sakthikumar, S., Anderson, M. Z., Berlin, A., Gujja, S., et al. (2015). Genetic and phenotypic intra-species variation in Candida albicans. Genome Res. doi:10.1101/gr.174623.114.

Hoang, D. T., Chernomor, O., von Haeseler, A., Minh, B. Q. & Vinh, L. S. UFBoot2: Improving the Ultrafast Bootstrap Approximation. Mol. Biol. Evol. 35, 518–522 (2018).

Holden DW. (1994) DNA mini prep method for Aspergillus fumigatus (and other filamentous fungi), p 3–4. In Maresca B, Kobayashi GS (ed), Molecular biology of pathogenic fungi, a laboratory manual. Telos Press, New York, NY.

Hong, S. B., Go, S. J., Shin, H. D., Frisvad, J. C., and Samson, R. A. (2005). Polyphasic taxonomy of Aspergillus fumigatus and related species. Mycologia. doi:10.3852/mycologia.97.6.1316.

Hu, W., Sillaots, S., Lemieux, S., Davison, J., Kauffman, S., Breton, A., et al. (2007). Essential gene identification and drug target prioritization in Aspergillus fumigatus. PLoS Pathog. doi:10.1371/journal.ppat.0030024.

Huson, D. H., and Weber, N. (2013). Microbial community analysis using MEGAN. Methods Enzymol. 531, 465–485.

Jiménez-Ortigosa, C., Moore, C., Denning, D. W., and Perlin, D. S. (2017). Emergence of echinocandin resistance due to a point mutation in the fks1 gene of Aspergillus fumigatus in a patient with chronic pulmonary aspergillosis. Antimicrob. Agents Chemother. doi:10.1128/AAC.01277-17.

Jones, P., Binns, D., Chang, H. Y., Fraser, M., Li, W., McAnulla, C., et al. (2014). InterProScan 5: Genome-scale protein function classification. Bioinformatics 30, 1236–1240.

Katoh, K., and Standley, D. M. (2013). MAFFT multiple sequence alignment software version 7: Improvements in performance and usability. Mol. Biol. Evol. doi:10.1093/molbev/mst010.

Katz, M. E., Dougall, A. M., Weeks, K., and Cheetham, B. F. (2005). Multiple genetically distinct groups revealed among clinical isolates identified as atypical Aspergillus fumigatus. J. Clin. Microbiol. doi:10.1128/JCM.43.2.551-555.2005.

Keller, N. P. (2017). Heterogeneity confounds establishment of “a” model microbial strain. MBio. doi:10.1128/mBio.00135-17.

Kjærbølling, I., Vesth, T. C., Frisvad, J. C., Nybo, J. L., Theobald, S., Kuo, A., et al. (2018). Linking secondary metabolites to gene clusters through genome sequencing of six diverse Aspergillus species. Proc. Natl. Acad. Sci. U. S. A. doi:10.1073/pnas.1715954115.

Klopfenstein, D. V., Zhang, L., Pedersen, B. S., Ramírez, F., Vesztrocy, A. W., Naldi, A., et al. (2018). GOATOOLS: A Python library for Gene Ontology analyses. Sci. Rep. doi:10.1038/s41598-018-28948-z.

Kowalski, C. H., Beattie, S. R., Fuller, K. K., McGurk, E. A., Tang, Y. W., Hohl, T. M., et al. (2016). Heterogeneity among isolates reveals that fitness in low oxygen correlates with Aspergillus fumigatus virulence. MBio. doi:10.1128/mBio.01515-16.

Kusuya, Y., Sakai, K., Kamei, K., Takahashi, H., and Yaguchi, T. (2016). Draft genome sequence of the pathogenic filamentous fungus Aspergillus lentulus IFM 54703T. Genome Announc. doi:10.1128/genomeA.01568-15.

Kusuya, Y., Takahashi-Nakaguchi, A., Takahashi, H., and Yaguchi, T. (2015). Draft genome sequence of the pathogenic filamentous fungus Aspergillus udagawae strain IFM 46973T. Genome Announc. doi:10.1128/genomeA.00834-15.

Lamoth, F. (2016). Aspergillus fumigatus-related species in clinical practice. Front. Microbiol. doi:10.3389/fmicb.2016.00683.

Latgé, J. P., and Chamilos, G. (2020). Aspergillus fumigatus and aspergillosis in 2019. Clin. Microbiol. Rev. doi:10.1128/CMR.00140-18.

Lê, S., Josse, J., and Husson, F. (2008). FactoMineR: An R package for multivariate analysis. J. Stat. Softw. doi:10.18637/jss.v025.i01.

Li, H., and Durbin, R. (2009). Fast and accurate short read alignment with Burrows-Wheeler transform. Bioinformatics. doi:10.1093/bioinformatics/btp324.

Li, H., Handsaker, B., Wysoker, A., Fennell, T., Ruan, J., Homer, N., et al. (2009). The Sequence Alignment/Map format and SAMtools. Bioinformatics 25, 2078–2079.

Lind, A. L., Wisecaver, J. H., Lameiras, C., Wiemann, P., Palmer, J. M., Keller, N. P., et al. (2017). Drivers of genetic diversity in secondary metabolic gene clusters within a fungal species. PLoS Biol. doi:10.1371/journal.pbio.2003583.

Liu, W., Sun, Y., Chen, W., Liu, W., Wan, Z., Bu, D., et al. (2012). The T788G mutation in the cyp51C gene confers voriconazole resistance in Aspergillus flavus causing aspergillosis. Antimicrob. Agents Chemother. doi:10.1128/AAC.05477-11.

McKenna, A., Hanna, M., Banks, E., Sivachenko, A., Cibulskis, K., Kernytsky, A., et al. (2010). The genome analysis toolkit: A MapReduce framework for analyzing next-generation DNA sequencing data. Genome Res. doi:10.1101/gr.107524.110.

Mondon, P., De Champs, C., Donadille, A., Ambroise-Thomas, P., and Grillot, R. (1996). Variation its virulence of Aspergillus fumigatus strains in a murine model of invasive pulmonary aspergillosis. J. Med. Microbiol. doi:10.1099/00222615-45-3-186.

Morris, S. M., Albrecht, U., Reiner, O., Eichele, G., and Yu-Lee, L. Y. (1998). The lissencephaly gene product Lis1, a protein involved in neuronal migration, interacts with a nuclear movement protein, NudC. Curr. Biol. doi:10.1016/S0960-9822(98)70232-5.

Negri, C. E., Gonçalves, S. S., Xafranski, H., Bergamasco, M. D., Aquino, V. R., Castro, P. T. O., et al. (2014). Cryptic and Rare Aspergillus species in Brazil: Prevalence in clinical samples and in Vitro susceptibility to Triazoles. J. Clin. Microbiol. doi:10.1128/JCM.01582-14.

Nguyen, L. T., Schmidt, H. A., Von Haeseler, A., and Minh, B. Q. (2015). IQ-TREE: A fast and effective stochastic algorithm for estimating maximum-likelihood phylogenies. Mol. Biol. Evol. doi:10.1093/molbev/msu300.

Nierman, W. C., Pain, A., Anderson, M. J., Wortman, J. R., Kim, H. S., Arroyo, J., et al. (2005). Genomic sequence of the pathogenic and allergenic filamentous fungus Aspergillus fumigatus. Nature. doi:10.1038/nature04332.

Patterson, T. F., Thompson, G. R., Denning, D. W., Fishman, J. A., Hadley, S., Herbrecht, R., et al. (2016). Practice guidelines for the diagnosis and management of aspergillosis: 2016 update by the infectious diseases society of America. Clin. Infect. Dis. doi:10.1093/cid/ciw326.

Posch, W., Blatzer, M., Wilflingseder, D., and Lass-Flörl, C. (2018). Aspergillus terreus: Novel lessons learned on amphotericin B resistance. Med. Mycol. doi:10.1093/mmy/myx119.

Puértolas-Balint, F., Rossen, J. W. A., Oliveira dos Santos, C., Chlebowicz, M. M. A., Raangs, E. C., van Putten, M. L., et al. (2019). Revealing the Virulence Potential of Clinical and Environmental Aspergillus fumigatus Isolates Using Whole-Genome Sequencing. Front. Microbiol. doi:10.3389/fmicb.2019.01970.

Ries, L. N. A., Steenwyk, J. L., De Castro, P. A., De Lima, P. B. A., Almeida, F., De Assis, L. J., et al. (2019). Nutritional heterogeneity among aspergillus fumigatus strains has consequences for virulence in a strain- And host-dependent manner. Front. Microbiol. doi:10.3389/fmicb.2019.00854.

Rivero-Menendez, O., Cuenca-Estrella, M., and Alastruey-Izquierdo, A. (2019). In vitro activity of olorofim (F901318) against clinical isolates of cryptic species of Aspergillus by EUCAST and CLSI methodologies. J. Antimicrob. Chemother. doi:10.1093/jac/dkz078.

Robbins, N., Caplan, T., and Cowen, L. E. (2017). Molecular Evolution of Antifungal Drug Resistance. Annu. Rev. Microbiol. doi:10.1146/annurev-micro-030117-020345.

Rokas A, Mead ME, Steenwyk JL, Oberlies NH, Goldman GH (2020) Evolving moldy murderers: Aspergillus section Fumigati as a model for studying the repeated evolution of fungal pathogenicity. PLoS Pathog 16(2): e1008315

Sharma, C., Kumar, R., Kumar, N., Masih, A., Gupta, D., and Chowdhary, A. (2018). Investigation of multiple resistance mechanisms in voriconazole-resistant aspergillus flavus clinical isolates from a chest hospital surveillance in Delhi, India. Antimicrob. Agents Chemother. doi:10.1128/AAC.01928-17.

Simão, F. A., Waterhouse, R. M., Ioannidis, P., Kriventseva, E. V., and Zdobnov, E. M. (2015). BUSCO: Assessing genome assembly and annotation completeness with single-copy orthologs. Bioinformatics. doi:10.1093/bioinformatics/btv351.

Stanke, M., Steinkamp, R., Waack, S., and Morgenstern, B. (2004). AUGUSTUS: A web server for gene finding in eukaryotes. Nucleic Acids Res. 32.

Steenwyk, J. L., Shen, X.-X., Lind, A. L., Goldman, G. H. & Rokas, A. A Robust Phylogenomic Time Tree for Biotechnologically and Medically Important Fungi in the Genera Aspergillus and Penicillium. MBio 10, (2019).

Sugui, J. A., Peterson, S. W., Figat, A., Hansen, B., Samson, R. A., Mellado, E., et al. (2014). Genetic relatedness versus biological compatibility between Aspergillus fumigatus and related species. J. Clin. Microbiol. doi:10.1128/JCM.01704-14.

Szalewski, D. A., Hinrichs, V. S., Zinniel, D. K., and Barletta, R. G. (2018). The pathogenicity of Aspergillus fumigatus, drug resistance, and nanoparticle delivery. Can. J. Microbiol. doi:10.1139/cjm-2017-0749.

Takahashi-Nakaguchi, A., Muraosa, Y., Hagiwara, D., Sakai, K., Toyotome, T., Watanabe, A., et al. (2015). Genome sequence comparison of Aspergillus fumigatus strains isolated from patients with pulmonary aspergilloma and chronic necrotizing pulmonary aspergillosis. Med. Mycol. doi:10.1093/mmy/myv003.

Tavaré, S. Some probabilistic and statistical problems in the analysis of DNA sequences. Lect. Math. life Sci. 17, 57–86 (1986).

Taylor, J. W., Jacobson, D. J., Kroken, S., Kasuga, T., Geiser, D. M., Hibbett, D. S., et al. (2000). Phylogenetic species recognition and species concepts in fungi. Fungal Genet. Biol. doi:10.1006/fgbi.2000.1228.

Tsai, H. F., Chang, Y. C., Washburn, R. G., Wheeler, M. H., and Kwon-Chung, K. J. (1998). The developmentally regulated alb1 gene of Aspergillus fumigatus: Its role in modulation of conidial morphology and virulence. J. Bacteriol. doi:10.1128/jb.180.12.3031-3038.1998.

Vinet, L. & Zhedanov, A. A ‘missing’ family of classical orthogonal polynomials. J. Phys. A Math. Theor. 44, 085201 (2011).

Walker, B. J., Abeel, T., Shea, T., Priest, M., Abouelliel, A., Sakthikumar, S., et al. (2014). Pilon: An integrated tool for comprehensive microbial variant detection and genome assembly improvement. PLoS One. doi:10.1371/journal.pone.0112963.

Waterhouse, A. M., Procter, J. B., Martin, D. M. A., Clamp, M., and Barton, G. J. (2009). Jalview Version 2-A multiple sequence alignment editor and analysis workbench. Bioinformatics. doi:10.1093/bioinformatics/btp033.

Waterhouse, R. M., Tegenfeldt, F., Li, J., Zdobnov, E. M. & Kriventseva, E. V. OrthoDB: a hierarchical catalog of animal, fungal and bacterial orthologs. Nucleic Acids Res. 41, D358–D365 (2013).

Winnenburg, R. (2006). PHI-base: a new database for pathogen host interactions. Nucleic Acids Res. doi:10.1093/nar/gkj047.

Yang, Z. Maximum likelihood phylogenetic estimation from DNA sequences with variable rates over sites: Approximate methods. J. Mol. Evol. 39, 306–314 (1994).

Yang, Z. Among-site rate variation and its impact on phylogenetic analyses. Trends Ecol. Evol. 11, 367–372 (1996).

Zakaria, A., Osman, M., Dabboussi, F., Rafei, R., Mallat, H., Papon, N., et al. (2020). Recent trends in the epidemiology, diagnosis, treatment, and mechanisms of resistance in clinical Aspergillus species: A general review with a special focus on the Middle Eastern and North African region. J. Infect. Public Health. doi:10.1016/j.jiph.2019.08.007.

Zbinden, A., Imhof, A., Wilhelm, M. J., Ruschitzka, F., Wild, P., Bloemberg, G. V., et al. (2012). Fatal outcome after heart transplantation caused by Aspergillus lentulus. Transpl. Infect. Dis. doi:10.1111/j.1399-3062.2012.00779.x.

Zheng, B., Yan, L., Liang, W., and Yang, Q. (2019). Paralogous Cyp51s mediate the differential sensitivity of Fusarium oxysporum to sterol demethylation inhibitors. Pest Manag. Sci. doi:10.1002/ps.5127.

