## Supplementary Figures for "Genomic and phenotypic heterogeneity of clinical isolates of the human pathogens *Aspergillus fumigatus, Aspergillus lentulus* and *Aspergillus fumigatiaffinis*"

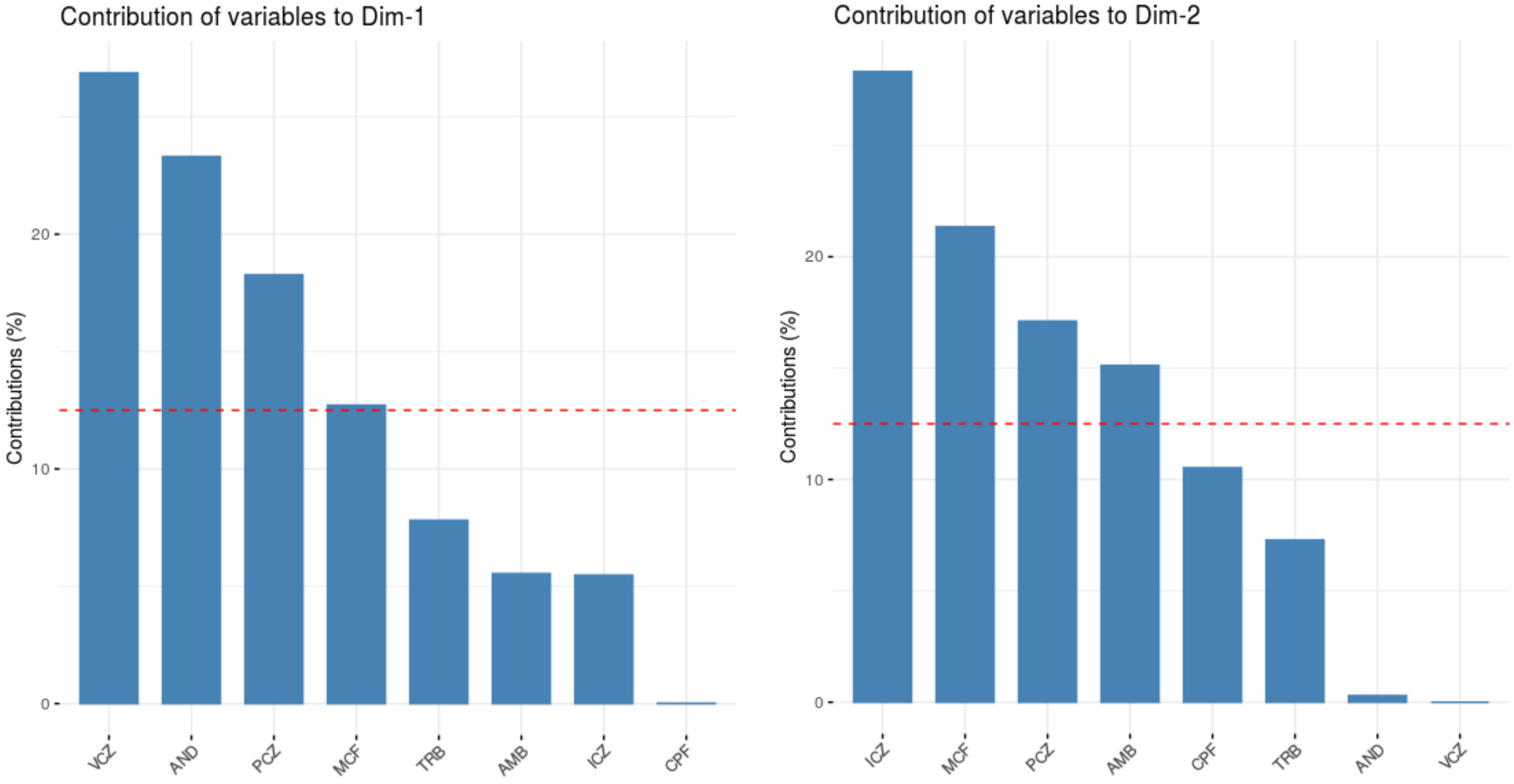


**Supplementary Figure 1**. Correlation between drug MIC/MEC and the principal component (PC).

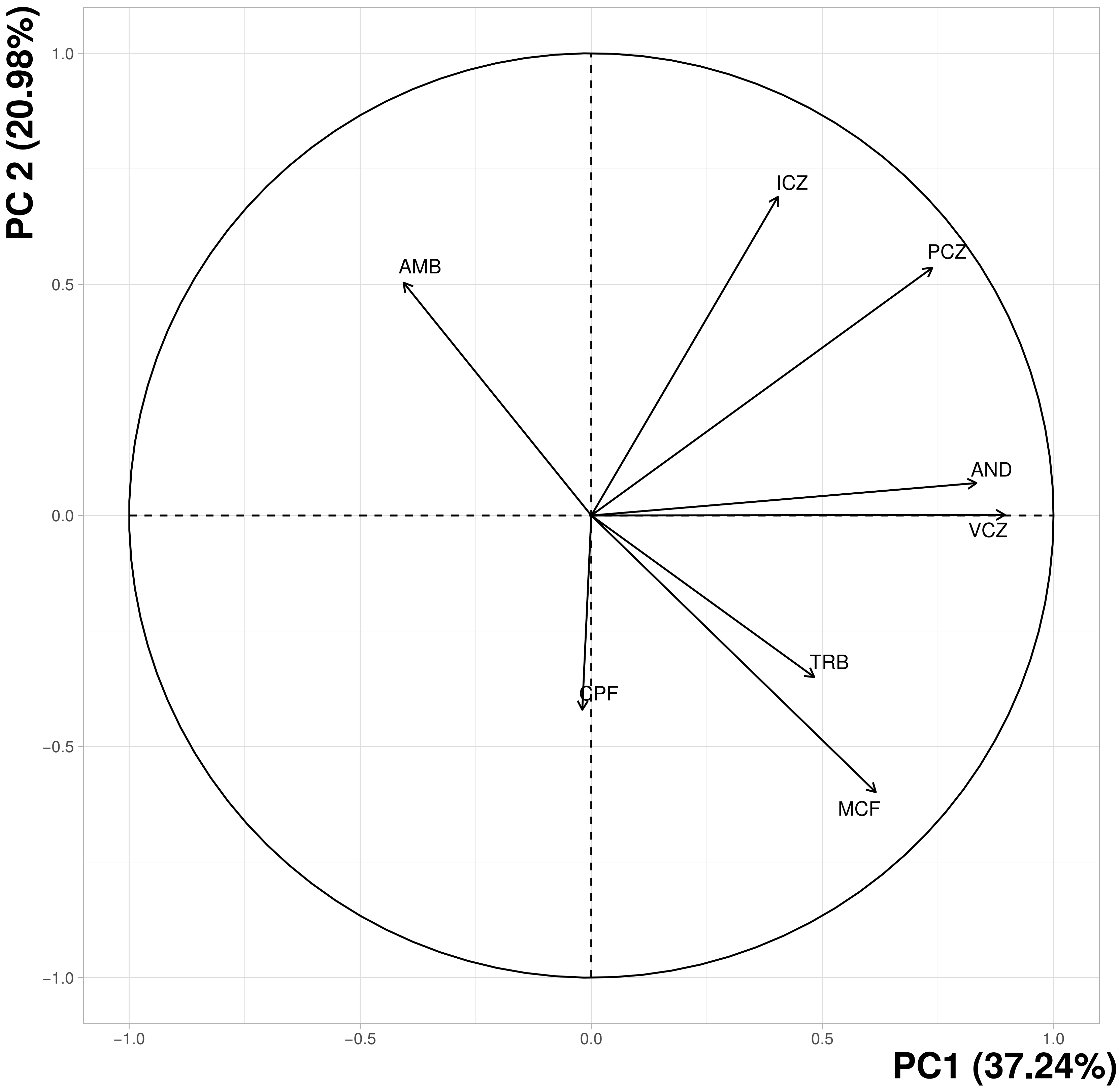


**Supplementary Figure 2**. Correlation circle with variables (antifungal MIC/MEC) contributing to each principal component (PC) in the PCA.

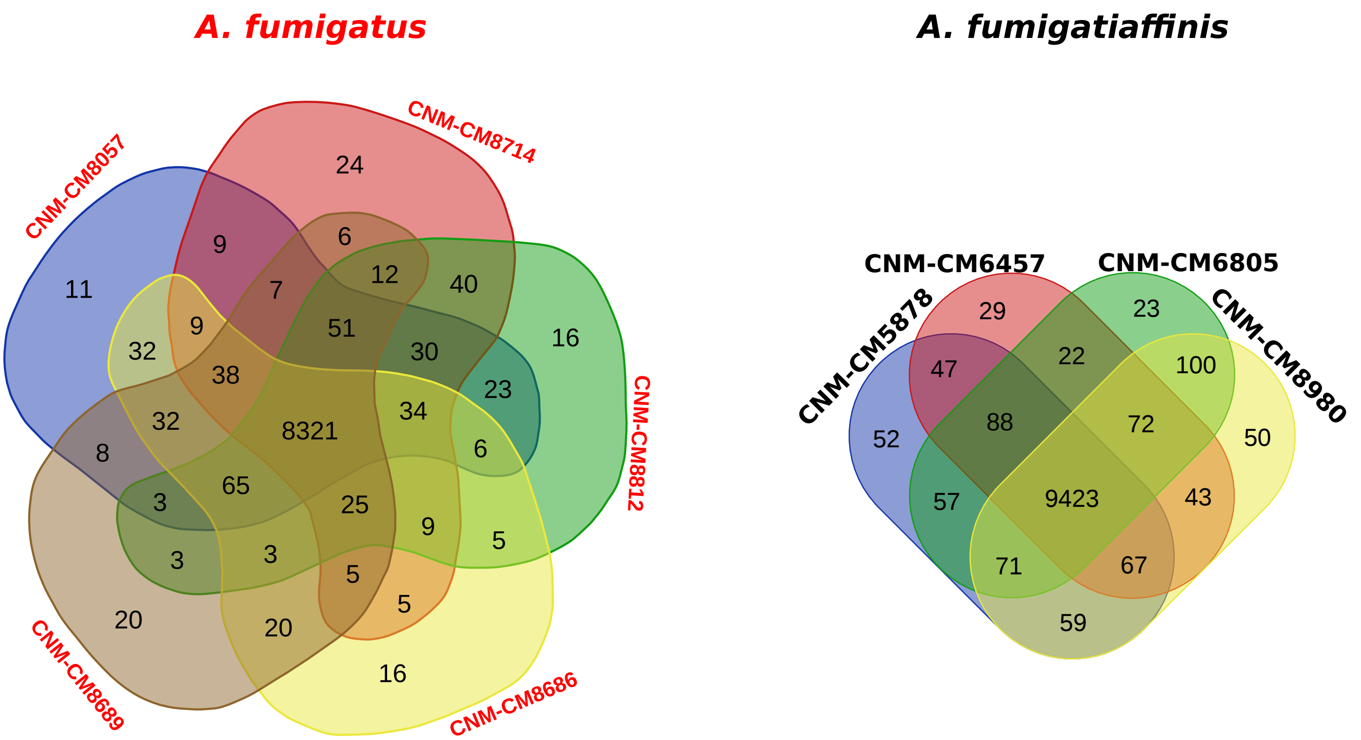


**Supplementary figure 3.** Venn Diagram of orthogroups shared by strains in each species for *A. fumigatus* and *A. fumigatiaffinis*.
